# Structure and function of the DFCP1 membrane barrier responsible for ER microdomain organization in lysosome repair

**DOI:** 10.64898/2026.10.19.759623

**Authors:** Zhicheng Cui, Karen Palacio-Rodriguez, Eric Herrmann, Athanasios Kournoutis, Maja Radulovic, Sascha Pust, Karla Tapia, Camilla Raiborg, Gerhard Hummer, Harald Stenmark, James H. Hurley

**Affiliations:** Department of Molecular and Cell Biology, University of California, Berkeley, CA, 94720, USA; California Institute for Quantitative Biosciences, University of California, Berkeley, CA, 94720, USA; Aligning Science Across Parkinson’s (ASAP) Collaborative Research Nework, Chevy Chase, MD 20815, USA; Department of Biochemistry and Molecular Genetics, University of Colorado Anschutz Medical School, Aurora, CO 80045, USA; Department of Theoretical Biophysics, Max Planck Institute of Biophysics, 60438 Frankfurt am Main, Germany; Center for Cancer Cell Reprogramming, Faculty of Medicine, University of Oslo, Montebello, Oslo, N-0424, Norway; Department of Molecular Cell Biology, Institute for Cancer Research, Oslo University Hospital, Montebello, Oslo, N-0379, Norway; Institute of Biophysics, Goethe University, 60438 Frankfurt am Main, Germany; Helen Wills Neuroscience Institute, University of California, Berkeley, Berkeley, CA 94720, USA

## Abstract

The endoplasmic reticulum (ER) forms specialized microdomains that organize autophagy, lipid droplet formation, and the repair of damaged lysosomes. The PI3P-binding protein DFCP1/ZFYVE1 marks these microdomains, raising the question as to whether and how one protein could define such a subcompartment. Here we determine the structure of full-length human DFCP1 bound to membranes by cryo-electron tomography and single-particle cryo-electron microscopy. DFCP1 assembles into a helical coat, built from tetramers that stack around a constricted membrane tube. Its ATPase domains form the organizing core of the helix, while its ER-targeting domains insert deeply into the outer leaflet, and its FYVE domains grip PI3P. Coarse-grained molecular dynamics simulations show that the coat does not simply follow the curvature of the membrane but acts as a barrier to lipid diffusion, almost completely blocking the movement of its cognate lipid PI3P past the filament and concentrating it beneath the coat. Point mutations that break the assembly, membrane-insertion or PI3P-binding interfaces each impair lysosomal repair in cells, and the assembly is also required for the focal recruitment of the lipid transporter VPS13C. We propose that DFCP1 collars fence off a specialized ER microdomain, so that locally made lipids can reach sites of repair rather than escaping into bulk ER.

---

The lysosome has been compared to the digestive tract of the cell because of its role in processing inbound material into useful metabolites^1^. Moreover, the lysosome is a central hub for metabolic signaling in the cell^2^. Over the past decade, it has become clear that lysosomes undergo active cycles of damage and repair, with repair orchestrated by an elaborate machinery^3,4^. This machinery includes the ESCRT membrane remodeling and resealing proteins^5,6^, the conjugation of ATG8s to single membranes (CASM) and LC3/GABARAP-assisted stimulator for ESCRT recruitment (LASER) pathways^7,8^, phosphatidylinositol 4-kinase 2A (PI4K2A)^9^, Oxysterol-binding protein related-proteins (ORPs)^9^, and the bridge-like lipid transporters (BLTPs) ATG2A^9^ and VPS13C^10^. The role of the Parkinson’s disease (PD) gene product VPS13C ^11^ connects dysfunction in lysosome repair to PD^10^. Lysosome repair events are localized to endoplasmic reticulum (ER)-lysosome membrane contact sites (MCS)^3,4^.

The complexity of the process begs the question as to what architectural principles organize ER-lysosome membrane contact sites (MCS) involved in lysosome repair. The lipid phosphatidylinositol 3-phosphate (PI3P) and the PI3P-binding protein DFCP1 were recently shown to be required for the organization of ER microdomains involved in lysosome repair^12^. Inhibition of PI3P production and knockout of DFCP1 were both shown to impair the recovery of lysosomal function following treatment by the damaging agent L-leucyl-L-leucine methyl ester (LLOMe)^12^. DFCP1 is required for the efficient recruitment of VPS13C to sites of repair, yet does not appear to directly interact with VPS13C in cells^12^. These observations suggest that DFCP1 is responsible in some way for organizing repair-mediating ER-lysosome MCS, yet how this might occur is unknown.

In an analogous role in macroautophagy, DFCP1 has long been known to mark a PI3P-enriched microdomain of the ER known as the “omegasome”, which is associated with autophagy initiation in mammalian cells^13^. The transfer of phospholipids by ATG2A at ER-phagophore MCS in autophagy appears to be functionally analogous to the role of BLTPs at ER-lysosome MCS in lysosome repair^14^. DFCP1 contains an ATPase domain related to that of atlastin and the GTPase domain of dynamin, but the other domains of DFCP1 are distinct. While DFCP1 is non-essential for bulk autophagy^13^, mutations that knock out the ATPase activity of DFCP1 impair its function in selective autophagy^15^. DFCP1 and PI3P are involved in lipid droplet (LD) biogenesis^16,17^, a process that depends on ER-LD contact sites in a manner that parallels lipid transfer in lysosome repair and autophagy. DFCP1 is also involved in the biogenesis of double membrane vesicles (DMV) that are derived from the ER ^18^. These observations suggest that DFCP1 could have a common PI3P-dependent function in ER microdomain organization.

DFCP1 consists of an N-terminal ATPase domain, a central ER-targeting domain (ERTD), and two C-terminal FYVE domains. The crystal structure of the isolated ATPase domain was determined, and it was shown to form a dimer stabilized in a molecular glue-like manner by ATP^19^. The ERTD is unrelated in sequence to any domain of known structure. The FYVE domains are PI3P-binding competent and are predicted to resemble the structures of other PI3P-binding FYVE domains^20,21^. It is unknown how these domains interact with each other in three dimensions or how they interact with membranes. We determined the structure of a membrane-bound assembly of full-length human DFCP1 by cryo-electron tomography (cryo-ET) and single particle cryo-electron microscopy (cryo-EM). As described below, the structure led to a testable hypothesis that helical collars of DFCP1 around tubular PI3P-containing membranes of ER-like composition can act as diffusion barriers, suggesting a mechanism for the compartmentalization of lipids and proteins in specialized ER microdomains.

## Results

To understand how DFCP1 might organize ER microdomain membranes, we purified full-length human DFCP1 and incubated it with large unilamellar vesicles (LUVs) composed of 84.8% DOPC, 10% cholesterol, 5% PI3P and 0.2% DiR. Cryo-electron tomography (cryo-ET) of the decorated liposomes revealed a heterogeneous population of remodeled membranes (Fig. 1a). On a subset of tubes, the protein formed an organized lattice on the membrane (Fig. 1a, red arrows). On others it formed a disordered, “fuzzy” coat that lacked obvious periodic order, yet still drove tubulation (Fig. 1a, blue arrows). We focused sub-tomogram averaging (STA) on the ordered tubes and obtained a map at 25 Å resolution (Fig. 1b, Extended Data Fig. 1a). The assembly has a periodicity of 17 nm and an inner membrane diameter of 5.5 nm (Fig. 1c). Both the STA map and the IsoNet2-denoised tomogram show a curved bilayer lying immediately beneath the protein coat (Fig. 1c,d, Extended Data Fig. 1c). We validated the map by template matching followed by back-projection into the original tomograms, which showed that the ordered lattice is prevalent across the constricted segments and that the DFCP1 coat is continuous along the tubes (Fig. 1e, Extended Data Fig. 1d).

**Figure 1.**
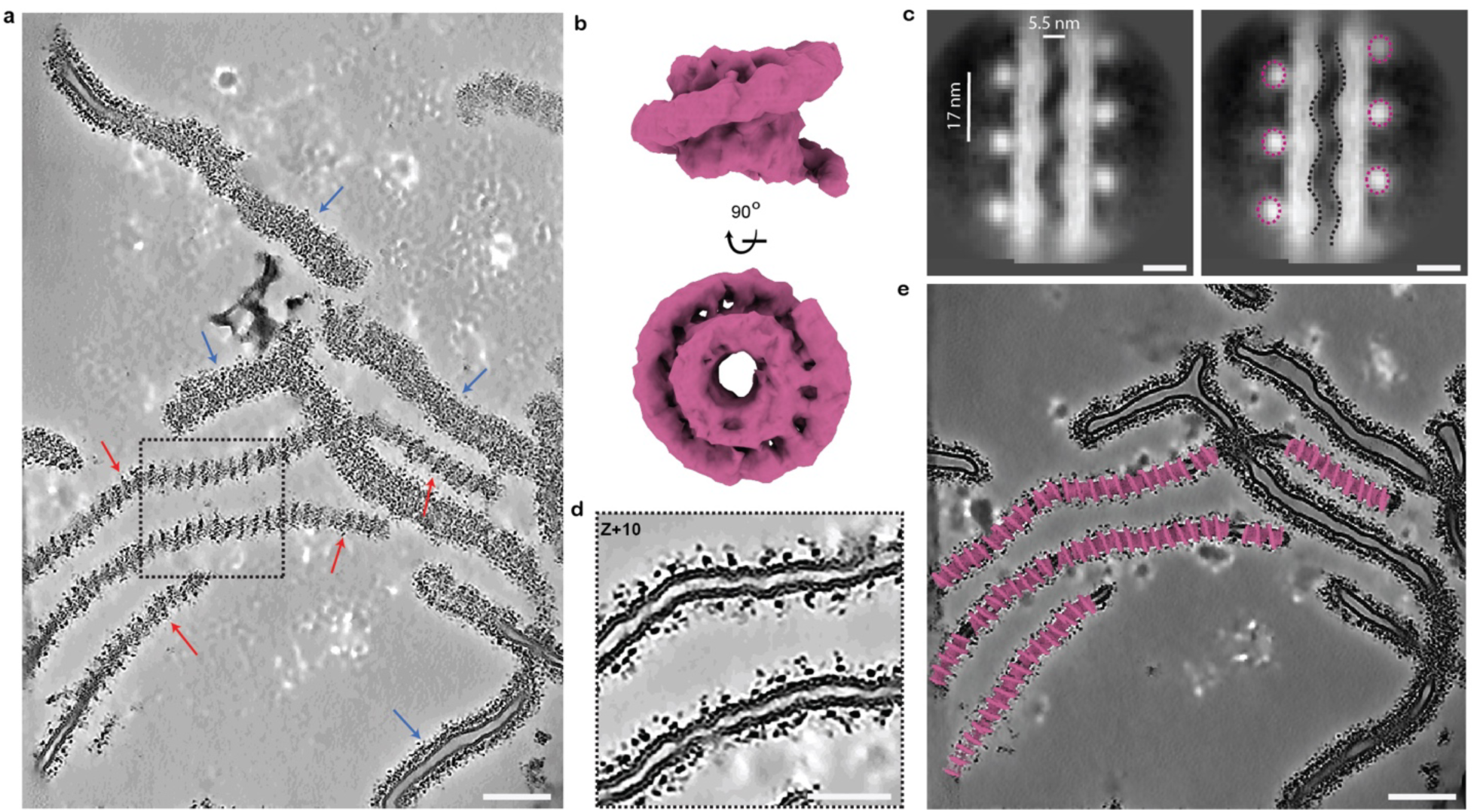
Architecture of DFCP1-decorated membrane tubes. **a**, A representative tomographic slice of full-length human DFCP1 reconstituted on liposomes (84.8% DOPC, 10% cholesterol, 5% PI3P, 0.2% DiR). Red arrows indicate the “constricted” coat morphology, while blue arrows highlight regions with a disordered, “fuzzy” coat morphology. Scale bar, 100 nm. **b**, Sub-tomogram averaging (STA) map of the constricted coat at 25 Å resolution, shown in top and side views. The average is focused on one repeating unit of the DFCP1 assembly (magenta surface). **c**, Close-up view of a constricted tube from (**a**). The inset highlights tight protein binding into the membrane bilayer that induces pronounced local curvature. Scale bar, 50 nm. **d**, 2D slice through the STA volume. The cross-section illustrates the tightly curved membrane (black) aligning precisely with the periodic positioning of the protein units (Purple). Scale bar, 10 nm. **e**, Validation of the map derived from STA. Visual assessment of the back-projected volume after template matching (TM) demonstrates a robust fit of the molecular model back into the original tomogram and highlights the continuous turning of DFCP1 along the membrane. Scale bar, 100 nm.

We next used single-particle cryo-electron microscopy (cryo-EM) to resolve the repeating unit of the coat at high resolution (Extended Data Fig. 2). The low-resolution cryo-ET data suggested a helical assembly, but the high-resolution 2D class averages showed that the arrays do not obey perfect helical symmetry, which ruled out conventional helical reconstruction. We therefore used the low-resolution STA volume as an initial model for helical refinement without imposing fixed helical parameters (Extended Data Fig. 2a). This allowed precise alignment of the particles and yielded a high-quality map that we used for subsequent refinement.

We obtained a map of five adjacent tetramers at 3.68 Å resolution (Fig. 2a). As in the cryo-ET data, the map resolved the curvature of the bilayer immediately beneath the protein units (Fig. 2b). Focused refinement on the tetrameric building block improved the resolution to 2.76 Å for the tetrameric ATPase domain with the ERTD and 3.07 Å for the monomeric ATPase domain with the ERTD and FYVE domains (Extended Data Fig. 2a). The PI3P headgroup bound the FYVE2 (Fig. 2h), nucleotides, side chains and zinc ions were all clearly resolved (Extended Data Fig. 3). The DFCP1 tetramer is a dimer-of-dimers with a wing-like architecture. Each protomer is built from three modules, a core ATPase domain, an ERTD, and two C-terminal FYVE domains. The tetramer is oriented so that all four pairs of FYVE domains and all four ERTDs face the membrane, forming a broad, multivalent platform that spans the tube (Fig. 2c).

**Figure 2.**
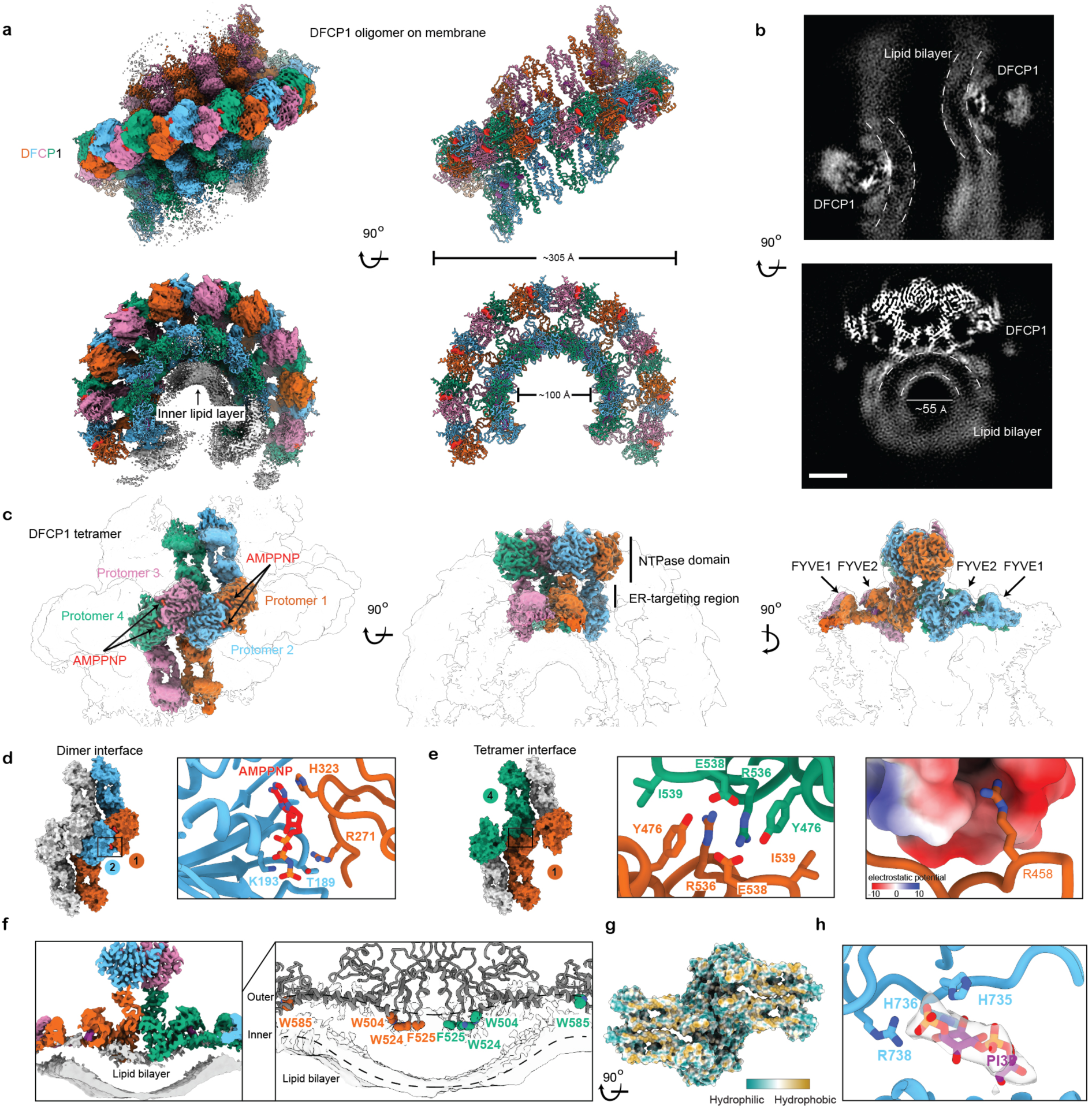
High-resolution cryo-EM structure of the membrane-bound DFCP1 tetramer. **a**, cryo-EM map and model of the higher-order assembly of DFCP1. Side and top views of the density map for the five-tetramer scaffold. The density clearly resolves the periodic protein units and the associated membrane. Dimensions of the assembly are indicated. **b**, Slice view of the cryo-EM map indicates pronounced local curvature indicated by the dashed lines. **c**, Composite map of the DFCP1 tetramer building block based on the focused maps and the overall map with C2 symmetry. Protomers 1-4 are colored orange-red, sky blue, reddish purple, and bluish green, respectively. Bound nucleotides (AMPPNP) are shown in red. Domain organization is labeled. **d**, Detailed view of the ATP-mediated primary dimer interface. The nucleotide is “sandwiched” between protomer 1 and 2. **e**, Structure of the novel tetramerization interface. Left: Surface view of the ERTDs of protomers 1 and 4, the ATPase domains of protomers 2 and 3 were omitted for better visualization. Center: Zoom-in view showing the hydrophobic stacking. Right: Surface electrostatic potential map highlighting the residue R458. **f**, Left: Slice through the composite tetramer map showing the protein model deeply embedded into the outer lipid layer, creating a local indentation that forces the inner lipid layer to curve, indicated by dashed lines. Right: Zoom-in view of the hydrophobic residues dipping into the space of acyl chain. **g**, Surface representation of DFCP1 tetramer colored by hydrophobicity, viewing from the membrane-interaction side. **h**, Detailed interactions in the PI3P-binding pocket with PI3P density.

Two interfaces hold the tetramer together. The first is an ATP-dependent dimer interface that matches the one seen in the crystal structure of the isolated ATPase domain^19^. It is formed by the symmetric association of the core ATPase domains of protomers 1 and 2, and of protomers 3 and 4 (Fig. 2c), and depends on the bound nucleotide, which is sandwiched between Lys193 and Thr189 of one protomer and His323 and Arg271 of the other (Fig. 2d). In essence, this protomer interface is held together by ATP in a molecular glue-like manner, as also seen previously^19^. The second is a tetramerization interface that we had not anticipated and that drives the higher-order assembly of the coat. It is formed by the ERTDs of protomers 1 and 4, which establish a second trans-protomer linkage within the tetramer (Fig. 2e). Here Tyr476, Arg536 and Ile539 form a hydrophobic stack at the core of the interface, reinforced by an electrostatic contact from Arg458 of the opposing protomer (Fig. 2e).

The tetramer is embedded deeply in the outer leaflet of the bilayer. It presses the outer leaflet inward, forcing the inner leaflet to follow and narrowing the lumen of the tube, therefore creating a saddle-shaped lipid bilayer under the DFCP1 oligomer (Fig. 2f, Extended Data Fig. 4a). Embedding is driven by an extensive hydrophobic surface, with Trp504, Trp524, Phe525, and Trp585 of each protomer inserting directly into the hydrophobic core of the membrane (Fig. 2f, g). Trp524 is the most deeply buried residue, roughly 10 Å below the plane of the phospholipid headgroups. Membrane anchoring is reinforced by multivalent recognition of PI3P by the FYVE domains. FYVE2 contacts the ATPase core and is better resolved than FYVE1, and its bound PI3P is clearly defined, with His736 and Arg738 forming a hydrogen bond and a salt bridge to the headgroup (Fig. 2h). The coat therefore stabilizes the tubular geometry of the membrane through a combination of deep hydrophobic indentation and high-affinity phosphoinositide binding.

We used coarse-grained molecular dynamics to test whether the coat resolved by cryo-EM is stable in a bilayer and how it impacts the coated membrane. We built Martini 3 models of the DFCP1 filament wrapped around a membrane nanotube of ER-like composition supplemented with 10% PI3P. As a control, we built a protein-free tube of matched radius, box size, and lipid composition (Fig. 3a,b). Accommodating the unusually deep penetration of the ERTD required removing a substantial fraction of the outer-leaflet lipids, on average about 30% of those under the footprint of each monomer (Extended Data Fig. 7). Over 15 µs of production in three independent replicates per system, the filament held a stable, curved conformation on the tube. The diameters of the protein ring (∼18.8 nm) and its membrane interface (∼11.5 nm) did not drift (Fig. 3d). The tube reached a stationary geometry within the first microsecond and kept outer- and inner-leaflet diameters close to those of the protein-free control in all replicate trajectories (Fig. 3e). The filament therefore neither pinches off nor destabilizes the tube it coats, in agreement with the coat architecture resolved by cryo-EM.

**Figure 3.**
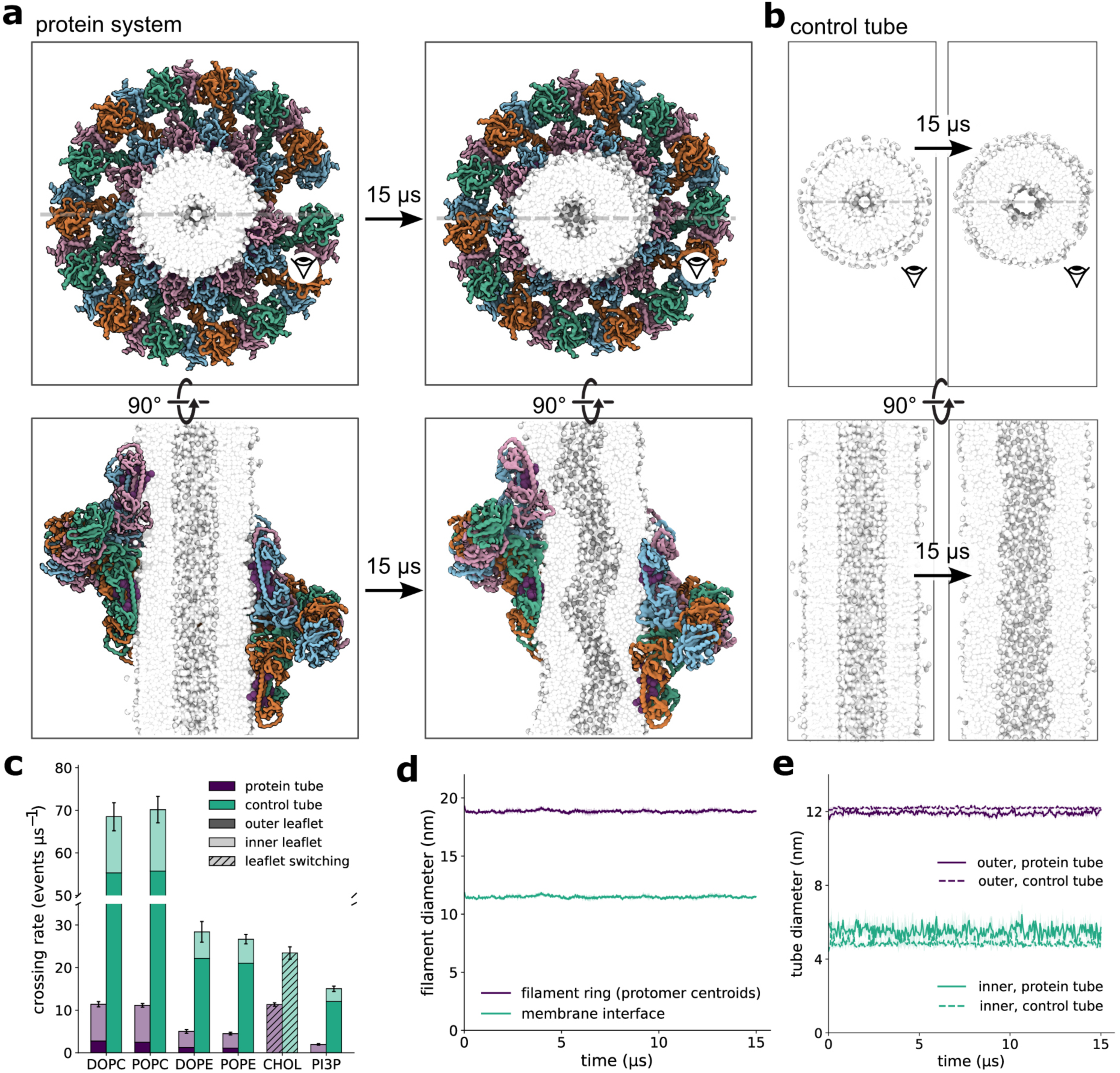
Molecular dynamics simulations of the DFCP1 filament on a membrane tube. **a,** Snapshots of the DFCP1 filament (protein system) on a membrane tube, at the start of the simulation (left) and after 15 µs of production (right, representative snapshots for one of three independent replicates). Top row, view along the tube axis; bottom row, cut-away view exposing the tube lumen. The four protomer chains that make up each of the filament’s seven tetramers are colored blue, green, orange and pink; PI3P lipids docked in the filament’s FYVE-domain pockets are shown in purple; lipid headgroups are shown in grey and lipid tails in white for every other lipid species. The solvent is not shown. **b,** As in **a**, for the protein-free control tube, which has the same tube radius, box dimensions and lipid composition as the protein system but lacks the filament and its docked PI3P. **c,** Axial crossing rate of each lipid species across the filament (protein system, purple) or, in the control tube (green), across a band of the same axial width placed at the tube’s center, reported as crossing events per µs of trajectory. Each bar stacks outer-leaflet crossings (dark shade), inner-leaflet crossings (light shade) and transits during which the lipid switched leaflet (hatched; only cholesterol, which crosses exclusively by flip-flop), so that its total height is the combined crossing rate; segments are means and error bars the s.d. of the total over n = 3 replicates per system. The y-axis is broken between 35 and 50 events µs⁻¹ to accommodate the control tube’s higher DOPC/POPC rates without compressing the other species; the leaflet-resolved breakdown is shown in Extended Data Fig. 5. **d,** Diameter of the filament measured about its own axis, as a function of time: the diameter of the cylinder fitted to the 28 protomer centroids (filament ring, dark purple) and twice the mean distance of the membrane-facing backbone beads from that axis (membrane interface, green), mean ± s.d., n = 3 replicates. **e,** Outer-(dark purple) and inner-leaflet (green) diameter of the membrane tube as a function of time, for the filament system (solid) and the control tube (dashed), tracked from the start of production (mean ± s.d., n = 3 replicates per system).

The simulations further show that the filament acts as a lipid diffusion barrier, and that the barrier is strongest for its cognate lipid, PI3P. In the outer leaflet, which the FYVE domains engage, PI3P crossed the band occupied by the filament only once in 45 µs of combined sampling, against 12.0 ± 0.7 crossings per µs on the control tube, an almost complete block (Fig. 3c). Summed over both leaflets, PI3P crossed at 13% of the control rate, the largest reduction of any species. The other phospholipids were held back almost as strongly, to between 16 and 18% of their control rates, and in every case the diffusion block was far stronger in the outer leaflet that the filament contacts than in the inner leaflet that it does not (Extended Data Fig. 5). Cholesterol was the only lipid to escape the fence, bypassing it by flipping into the inner leaflet, and even this route ran at just under half the control rate. The strong block of phospholipid diffusion acts on top of the entropic constriction effect, which reduces lipid flux through a pore constriction as ∼1/R with R the radius of the constriction. The barrier also concentrates PI3P where the filament binds it. Free PI3P, beyond the molecules already docked in the FYVE pockets, was recruited into the outer leaflet directly beneath the filament, with no comparable recruitment in the inner leaflet (Extended Data Fig. 6).

Together these results indicate that DFCP1 is not a passive coat that conforms to a pre-existing curvature, but an active organizer of the bilayer. It holds a stable curved geometry over microsecond timescales while sequestering its cognate lipid, PI3P, in the leaflet it contacts. This combination of stable coat geometry, lipid sequestration, and reduced lipid diffusion, which is inaccessible to structural methods alone, provides a mechanistic basis for how a curvature-stabilizing filament could compartmentalize a lipid subdomain on a membrane tube in vivo. A fuller account of the simulations is given in the Extended Analysis of MD Simulations.

DFCP1 is required for efficient repair of damaged lysosomes, and this depends on its ability to bind and hydrolyze ATP ^12^. To ask whether the ordered assembly we resolved is also required, we mapped point mutations onto the three interfaces that build the coat and tested them in cells. We stably expressed mCherry-tagged WT DFCP1 or one of three mutants in U2OS cells lacking endogenous DFCP1, a tetramer-interface mutant (R458D/R536D/I539D), a membrane-embedding mutant (W504D/W524D/F525D/W585D) and a PI3P-binding mutant (H618D/R621E/H735D/R738E), and followed the recovery of lysosomal integrity after LLOMe damage with the LysoTracker assay^6^. Cells expressing WT DFCP1 restored lysosomal integrity, whereas repair was strongly impaired by all three mutants (Fig. 4a). The interface that holds the coat together, the surface that embeds it in the membrane and the pocket that binds PI3P are therefore each required for DFCP1 to function in lysosomal repair. These results indicate that ordered assembly of DFCP1 on ER membranes is required for its role in repair.

**Figure 4.**
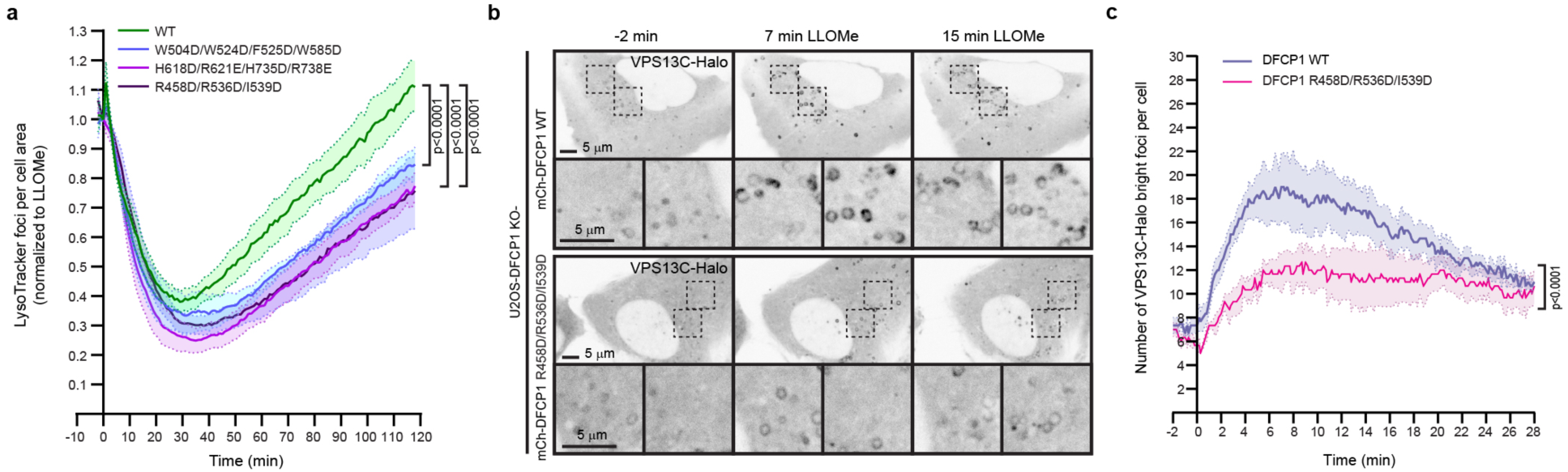
Role of structural interfaces in lysosomal membrane repair and VPS13C localization. **a,** U2OS DFCP1 KO cells stably transfected with mCherry-tagged versions of WT DFCP1 or the membrane-bound tetramer interface (R458D/R536D/I539D) mutant, the ERTD membrane embedding (W504D/W524D/F525D/W585D) mutant or the PI3P binding (H618D/R621E/H735D/R738E) mutant were pretreated with 75 nM LysoTracker Deep Red for 30 min before the addition of 250 μM LLOMe. Cells were analyzed by live spinning-disk microscopy (1 frame per minute for 120 minutes), and LysoTracker fluorescent bright puncta per cell area was measured. The graphs show mean +/-SEM from n=5 independent experiments, >200 cells per condition. The ordinary two-way ANOVA, Tukey’s multiple comparison test was used. Source numerical data are available in source data. **b,** Representative fluorescence micrographs of U2OS DFCP1 KO cells stably transfected with mCherry-tagged WT DFCP1 or the DFCP1 membrane-bound tetramer interface mutant and transiently expressing VPS13C-Halo following treatment with 250 µM LLOMe. In wild-type cells, VPS13C accumulated in discrete punctate microdomains whereas VPS13C recruitment appeared much more diffuse in cells expressing the tetramer mutant. Images were acquired every 10 s for 30 min. Scale bars: 5 µm. **c,** The quantification graph of VPS13C-Halo microdomain formation shown in b). LLOMe was added at time 0. Error bars denote mean ± SEM from n=3 independent experiments, >60 cells per condition were analyzed using ordinary two-way ANOVA.

Focal accumulation of the bridge-like lipid transporter VPS13C at damaged lysosomes depends on DFCP1^12^. To test whether the assembly, rather than DFCP1 alone, is what concentrates VPS13C, we followed Halo-tagged VPS13C by live spinning-disk microscopy in DFCP1-KO cells expressing WT DFCP1 or the tetramer-interface mutant. In cells with WT DFCP1, VPS13C rapidly accumulated on lysosomes after addition of LLOMe and gathered into hotspots that correspond to sites of membrane damage (Fig. 4b, upper panels), as found previously^12^. In cells expressing the tetramer-interface mutant, this focal accumulation was abolished and VPS13C remained diffuse (Fig. 4b, lower panels), a difference confirmed by quantification of the movies (Fig. 4c). The ordered DFCP1 coat is therefore needed to fence VPS13C into the focal sites that contact damaged lysosomes, as expected if the coat acts as a diffusion barrier.

## Discussion

By integrating cryo-ET and cryo-EM, we were able to visualize how the ATPase, ER targeting, and FYVE domains of DFCP1 are organized into a helical assembly with an inexact repeat. The assembly is organized hierarchically in that DFCP1 monomers are organized into tetramers, and the tetramers in turn serve as the protomers of the helical cable. The ATPase domains project away from the membrane. They serve as the main organizing element of the helix. The dimeric ATPase element of the tetramer matches that seen in the crystal structure of the isolated fragment ^19^. The dimer is held together by a bridge between the Arg271 side-chain of one unit with the γ-phosphate of ATP bound to the other, as also seen in the crystal structure ^19^. This explains why the mutants that are unable to hydrolyze ATP led to impaired turnover of the assembly as manifested in the loss of fluorescence recovery after photobleaching (FRAP) ^12^. This also shows how the dynamic turnover of the assembly by dissolution of the ATP-dependent protomer interface following ATP hydrolysis controls the lifetime of the diffusion barrier.

Although the ATPase domain itself has homology with the GTPase domain of dynamin, the structure of the helical assembly has no detailed similarity to those of assemblies of dynamin or dynamin-related proteins. DFCP1 contains no counterpart of the bundle signaling element (BSE) responsible for mechanotransduction from the GTPase domain to the membrane coat^22^. As compared to dynamin and its relatives, and indeed essentially all other helical membrane coats that we are aware of, the interactions between protein layers in successive turns are much more limited for DFCP1.

While there is no contact between the ATPase cores in successive helical turns, the ER targeting domain and FYVE1 project along the helical axis, pointing away from the central cable. The protrusion of the ER targeting domain and FYVE1 and the need to avoid steric clash sets the minimum spacing between successive helical turns. The distal tips of the ER targeting domain and FYVE1 from successive turns do not appear to form strong, regular interactions with one another. They seem to serve as spacer elements which set the minimum distance between successive helical turns in order to avoid steric overlap.

The degree of membrane penetration by the ER targeting domain is exceptional for a protein that does not span the membrane bilayer and is not classified as an integral membrane protein. The combination of deep membrane penetration with close approach of the tips of the ERTDs suggested to us that the ER targeting domain in the helical context could form a nearly impenetrable barrier to the diffusion of lipids in the outer leaflet. The results of MD simulations show that DFCP1 stably inserts into the membrane and forms a potent diffusion barrier that segregates lipids. DFCP1 immobilizes and concentrates its cognate lipid, PI3P, in the leaflet it contacts. Except for cholesterol, which can bypass the fence by flipping into the inner leaflet, all of the other lipids included in the simulation were virtually confined on one side of the barrier. The simulation only considered a single turn of the helix with a single close tip-to-tip approach forming a tight barrier. Multiple DFCP1 rings should further enhance the lipid confinement effect, acting in concert with the entropic effect of constriction. While membrane proteins were not included in the simulation, it is reasonable to assume that any barrier to diffusion of outer leaflet phospholipids will also completely block lateral diffusion of any membrane-associated proteins such as VPS13C.

The ER consists of a dense matrix of dynamic membrane tubules ^23^, whose cylindrical curvature is stabilized by reticulons and REEPs ^24^. The 130-amino acid DFCP1 ERTD is completely distinct architecturally from the 30-35 amino acid hairpin domains of the reticulons. Reticulons and REEPs do not contain nucleotide hydrolyzing domains, nor are they known to form helical structures such as the one reported here. The entirely distinct structures of DFCP1 as compared to reticulons and REEPs are in line with their entirely distinct biological functions. Reticulons and REEPs are the core determinants of ER tubular curvature. We propose that DFCP1 does not exist primarily to induce curvature, but rather to transiently segregate a specialized ER microdomain from bulk ER for the duration of a process such as lysosome repair or autophagy, occurring on a time scale of minutes. Given the matrix nature of ER tubules and their ability to dynamically branch and fuse, it is likely that several, and perhaps many, DFCP1 collars would be needed to sustain microdomain segregation over the distance scale of perhaps 100-300 nm and time scale of 10-30 minutes involved in lysosome repair and autophagy.

One of the unsolved puzzles in the BLTP field is what governs the directionality of phospholipid transport ^14^. BLTPs are passive transporters, uncoupled from an energy source. They allow phospholipids to move down a thermodynamic gradient across aqueous barriers, but the nature of the thermodynamic driving force is unknown. Local phospholipid synthesis in the source membrane (in this case, the ER) is an attractive candidate to establish such a driving force. Local synthesis of phospholipids is known to occur in the ER at sites of autophagosome biogenesis ^25^. It has been unclear, however, what would prevent any locally synthesized phospholipids from diffusing laterally into bulk ER. A lipid diffusion barrier contributed by DFCP1 would offer a solution, confining newly made phospholipids in a diffusionally restricted microdomain from which the only egress would be through the BLTP channel.

## Methods

### Protein expression and purification

Codon-optimized human full-length DFCP1 was synthesized (Twist Bioscience) and cloned into a pCAG vector with a tobacco etch virus (TEV)-cleavable MBP tag at the N terminus. For expression and purification of MBP-DFCP1, HEK293F GnTI^−^ cells were cultured in FreeStyle 293 Expression Medium (Thermo Fisher Scientific) and transfected at a density of 1.5-1.8 x 10^^6^ cells per ml with 1 mg of plasmid DNA AND 4 mg polyethylenimine (PEI) per liter. After 48-72 h of expression, cells were harvested by centrifugation at 800g for 20 min and resuspended in wash buffer (50 mM HEPES pH 7.5, 300 mM NaCl, 1 mM MgCl_2_, 1 mM TCEP, 1% Triton X-100) supplemented with 10% glycerol, and a protease inhibitor cocktail (Roche). Cell lysate was cleared by centrifugation at 35,000g for 35 min. The supernatant was incubated with Amylose resin (New England Biolabs) for 2 hr. at 4°C. The resin was first washed with wash buffer and then washed with wash buffer without the Triton X-100. The protein was eluted with 10 mM maltose and incubated with ATP (0.1 μM) overnight at 4°C. The MBP-tagged DFCP1 was further purified by size-exclusion chromatography using a Superdex 200 Increase 10/300 GL column (GE Healthcare) equilibrated in SEC buffer (50 mM HEPES pH 7.5, 150 mM NaCl, 1 mM MgCl_2_, 1 mM TCEP). Peak fractions corresponding to the monomeric MBP-DFCP1 were pooled, concentrated to ∼5.4 mg/ml, and immediately flash frozen in liquid nitrogen and stored in −80 °C.

### Large unilamellar vesicle preparation

Large unilamellar vesicles (LUVs) were prepared by mixing DOPC (Avanti Polar Lipids: 850375C), cholesterol (Avanti Polar Lipids: 700000P), PI3P (Avanti Polar Lipids: 850150), and DiR (Invitrogen: D12731) in a molar ratio of 84.8 : 10 : 5 : 0.2. The lipid mixture in chloroform was dried under a gentle nitrogen stream and desiccated overnight. The resulting lipid film was rehydrated in SEC buffer to a final concentration of 1.8 mM and subjected to 10 freeze-thaw cycles. LUVs were then generated by extruding the lipid suspension 61 times through a 30-nm polycarbonate membrane (Whatman Nuclepore Track-Etched Membranes, diameter of 19 mm) using a mini-extruder (Avanti Polar Lipids).

### DFCP1 assembly on membranes

For in vitro reconstitution, a total volume of 50 μL was used. Purified MBP-DFCP1 was incubated with LUVs at a concentration of 19 μM and 0.4 mM, respectively, for 2 h at room temperature in the presence of 2 mM AMPPNP and 5 mM MgCl_2_. The mixture was then centrifuged at 21,130xg to pellet the liposomes and resuspended in 20 μL of SEC buffer with 2 mM AMPPNP and 5 mM MgCl_2._ The resuspended liposomes were used for vitrification.

### Cryo-ET data collection and processing

3 μl of the sample was applied to a glow-discharged (PELCO easiGlow, 45 s in air at 15 mA and 0.37 mbar) holey carbon grid (C-flat, 2/1-3C-T) and vitrified using the FEI Vitrobot Mark IV System (Thermo Fisher Scientific). The samples were incubated on grids for 1 min and blotted for 3 s with a blot force of 15, using two Whatman 595 papers on the sample side and one Whatman 595 paper on the backside, at 6 °C with 95% relative humidity. Tilt series were collected on a Titan Krios (Thermo Fisher Scientific) operating at 300 kV using a K3 direct electron detector (Gatan) using super-resolution correlated double sampling (CDS) mode and BioQuantum energy filter (Gatan) with a 20 eV slit. Data were acquired using SerialEM ^26^ in a dose-symmetric scheme from ࢤ60° to +60° with 3° increments. The total dose was ∼120 e-/Å² and the pixel size was 0.9 Å. Scipion3 was used to facilitate downstream data processing. Images were motion-corrected using MotionCor3 (https://github.com/czimaginginstitute/MotionCor3; RRID:SCR_01649) ^27^, utilizing the anisotropic correction algorithm originally described before. While correction, the tilt series was binning to a pixel size of 3.62 Å and CTFfind5 ^28^ was used for CTF estimation. IMOD ^29^ was used for coarse prealinment, generation and the alignment of the gold fiducial model. Fidder was used to detect and erase the gold fiducials, before a final tomogram reconstruction with AreTomo2 ^30^ to a final pixel size of 10.85 Å. Tomograms were denoised and missing wedge corrected in two steps using IsoNet2 ^31^. The network training and volume post-processing were executed with a final B-factor of 200 A^2^ and a missing-wedge (MW) weighting factor of 200. A final denoising was performed using TOMO3D ^32^ to enhance visual contrast.

Particles were picked using Napari ^33^ and further processed with Relion5 ^34^. After creation of initial model another round of particle picking was performed using GAPSTOP ^35^. After processing Tomograms were visualized using 3DMOD ^36^ and the ChimeraX ^37^ based software ArtiaX ^38^.

### Single-particle cryo-EM data collection and processing

The same grids were used for cryo-EM data collection. Automated data acquisition was performed using SerialEM on a K3 Summit direct detection camera (Gatan) in super-resolution CDS mode with a pixel size of 0.52 Å and a defocus range of −0.9 to −2.2 μm. A total of maximum 245 exposures per stage shift were enabled by large beam shift. Beam intensity was adjusted to a dose rate of around 1 e^−^ Å^−2^ per frame for a 50-frame movie stack with a total exposure time of 7.1 s. A total of 21,780 movies were recorded.

Data processing was performed entirely within cryoSPARC (v.4) ^39^. A total of 21,780 micrographs were collected and subjected to Patch Motion Correction and Patch CTF Estimation. Micrographs were filtered based on defocus cutoff of 4 μm and a CTF resolution cutoff of 4 Å, resulting in a final set of 20,758 micrographs for processing. Particles were initially picked using blob picker from micrographs that were divided into 4 subsets. The initial particle stack from each subset was subjected to several rounds of 2D classification to only include the tubular structures. After combining the particles and removing duplicates, a clean set of 1,068,811 particles representing heterogeneous filamentous arrays was selected for 3D processing. For initial 3D reconstruction, the 25 Å resolution STA map was used as an initial model. To address the lack of perfect helical symmetry, helical refinement was conducted without the imposition of helical constraints. This “no-helical” refinement strategy enabled the accurate alignment of individual subunits within the half turn of the flexible filamentous arrays, yielding a reconstruction of a higher-order assembly at 4.12-Å resolution. This map was subsequently used for 3D classification and heterogeneous refinement to further clean the particles, and a non-uniform refinement job to produce a 3.74-Å map.

To resolve the higher-order structural heterogeneity, 825,653 particles were subjected to additional round of 3D classification. Three classes containing 589,296 particles showed the most well-defined features of the five-tetramer scaffold and were used to yield a 3.68-Å map. To achieve the highest possible resolution of the fundamental building block, a mask focused on the octamer around the central units was used for local refinement and then particle reextraction with a boxsize of 320. The octamer was further refined with a C2 symmetry and then symmetry-expanded to yield a 2.86-Å map of the tetramer. Particle subtraction was then performed in cryoSPARC to remove the contributions from adjacent tetramers and the underlying membrane. The resulting 1,194,950 subtracted symmetry-expanded particles were then used for local refinement of the tetramer with a C2 symmetry. The subtracted symmetry-expanded particles were then symmetry-expanded again to generate 2,389,900 particles that are focused on the tetramer. The final focused 3D classification was performed on these particles to select homogeneous tetrameric ATPase domain and the dimeric FYVE domain at C terminus. The final 3D reconstruction was performed using a soft-edged mask, yielding maps of tetrameric ATPase domain with ER-targeting region and the monomeric ATPase domain with ER targeting region and FYVE domain with a global resolution of 2.76 Å and 3.07 Å, respectively, according to the gold-standard Fourier shell correlation (FSC) 0.143 criterion. Local resolution was estimated using the Blocres algorithm in cryoSPARC.

### Atomic model building and refinement

The initial model of tetrameric DFCP1 was generated using AlphaFold2 ^40^ and rigid-body fitted into the cryo-EM map using UCSF ChimeraX. Manual model building and lipid docking (PI3P and structural lipids) were performed in Coot (v.0.9) ^41^. To optimize the protein-lipid and protein-nucleotide interfaces, interactive molecular dynamics flexible fitting was conducted using ISOLDE ^42^ within ChimeraX. The model was further refined using real-space refinement in PHENIX (v.2.0) ^43^ and validated using MolProbity ^44^ and EMRinger ^45^.

### Molecular Dynamics simulations

All simulations were performed with the Martini 3.0.0 coarse-grained force field ^46^, using GROMACS 2025.0 ^47^. The workflow comprised atomistic filament preparation, coarse-graining, membrane-tube construction, phosphoinositide docking, insertion of a membrane-embedded porin, solvation, ionization, equilibration, and production. Two systems were simulated: the DFCP1 filament wrapped around a membrane tube and a protein-free, curvature-matched membrane tube control. Identical simulation parameters, box geometry, tube radius, and lipid composition were used for both systems unless noted otherwise.

#### Filament structure preparation

The DFCP1 filament model comprised 28 protomers arranged as seven stacked tetramers around the tube axis. Residues Glu137–Leu777 of each protomer (641 residues per chain) were retained from the full-length model. The atomistic structure was energy-minimized (steepest descent) using the CHARMM36m ^48^ force field with the TIP3P water model ^49^, solvated in explicit water and 150 mM NaCl.

#### Coarse-graining

The minimized atomistic filament was converted to a Martini 3 representation with martinize2 (vermouth 0.14.0) ^50^. Secondary structure was assigned with DSSP ^51^. An elastic network was applied across all 28 chains (force constant 700 kJ mol⁻¹ nm⁻², lower/upper elastic bond cutoffs 0.5/0.9 nm), coupling beads between as well as within chains so that the filament’s inter-protomer contacts were retained through the tetramer interfaces.

#### Membrane tube construction

The membrane comprised DOPC (30%), POPC (30%), DOPE (12%), POPE (12%), the phosphoinositide PI3P (10%), and cholesterol (6%). A flat bilayer of this composition was first built with ‘insane.py’ ^52^ on a 20 × 20 × 12 nm template and, after removing the initial solvent, wrapped into a cylindrical membrane with BUMPy ^53^ (cylinder radius ≈ 5.9 nm), yielding a periodic tube of ∼64 nm length in a 38 × 38 × 64 nm box. The tube was built with its outer leaflet 2.0 nm smaller in radius than the filament’s PI3P-pocket circle (7.69 nm), giving a tube geometry (outer diameter ≈ 11.4 nm, lumen diameter ≈ 4.4 nm) that was used for both the protein system and the control tube. On this curved geometry the outer leaflet holds more lipids than the inner leaflet because its circumference is larger (outer:inner ≈ 1.5:1 by lipid count in both systems). Cholesterol was distributed the same way as every other lipid species during construction; during production it partitioned toward the inner leaflet, comprising 4.3% of outer-leaflet and 8.0% of inner-leaflet lipids in the protein system (4.6% and 7.7% in the control tube), with constant mole fractions over the 15 µs (Supplementary Fig. 3). The near-identical partitioning in both systems indicates that it is set by the tube curvature rather than by the filament.

#### PI3P docking

Fifty-six PI3P molecules were placed at paired histidine/arginine pockets on the filament (two sites per protomer, 28 protomers), by rigid-body superposition of a pre-equilibrated PI3P pose onto each protomer’s local frame. During energy minimization and equilibration, each docked PI3P was tethered to its pocket with a flat-bottom potential (lower bound 0 nm, upper bounds 0.3/0.6 nm, force constant 250 kJ mol⁻¹ nm⁻²), which was released for production.

#### Membrane-embedded porin

A single Martini 3 carbon-nanotube porin (5.0 nm diameter, 4.6 nm length, regular SC5 and functionalized end-ring SN2 beads) was generated with ‘martini-cnt-generator.py’ (bio-phys/cnt-martini) ^54,55^ and inserted radially through the tube wall, azimuthally and axially distant from the filament. Under the periodic, semi-isotropically coupled boundary conditions used here, the tube lumen would otherwise be a sealed solvent compartment; the porin allows the lumen to exchange solvent with the exterior compartment, so that the internal and external pressures remain physically coupled.

#### Solvation and ionization

All components were merged and any lipid within 0.4 nm of the filament, docked PI3P, or porin was removed. This clash-based removal is what carves the filament’s footprint into the membrane: in the protein system, where the filament and its 56 docked PI3P were present, it removed 420 lipids and cholesterol, predominantly from the outer leaflet beneath the filament; the same procedure applied to the same starting tube built without the filament removed 26 lipids clashing with the porin or other lipids in the control tube. The system was solvated with Martini water, water beads within 0.45 nm of membrane acyl-chain beads were removed, and the system was neutralized and ionized to 150 mM NaCl.

#### Simulation parameters, equilibration, and production

Non-bonded interactions used the Verlet cutoff scheme with a neighbor list updated every 20 steps. Electrostatic interactions used the reaction-field method (ε_r_ = 15, ε_rf_ = 0) ^56^, and both electrostatic and van der Waals interactions were truncated at 1.1 nm. Bond constraints were handled with LINCS ^57^. Energy minimization used steepest descent. Systems were equilibrated for 2 ns in the NVT ensemble at 310 K with the velocity-rescaling thermostat ^58^ (τ_t_ = 1.0 ps), followed by 10 ns in the NPT ensemble at 310 K and 1.0 bar with the Berendsen barostat ^59^ (semi-isotropic coupling, τ_p_ = 12 ps), under position restraints on the filament (and, for the protein-filament system, the flat-bottom PI3P restraints described above). Production runs used a 20 fs timestep, the velocity-rescaling thermostat (310 K, τ_t_ = 1.0 ps), and the C-rescale barostat ^60^ (semi-isotropic, 1.0 bar, τ_p_ = 12 ps), with the tube’s lateral (cross-sectional) compressibility set to zero, so that only the tube length along the periodic axis was free to relax and the tube radius could respond dynamically to the filament. All restraints were released for production. Three independent replicates of 15 µs were run for each system, with coordinates written every 1 ns.

#### Protein-free tube control

The control tube was built from the membrane tube used for the protein system, including the porin, and taken at the stage before filament insertion, so that the clash-based carving described above (420 lipids and cholesterol removed in the protein system versus 26 in the control tube) left the control tube with more lipids than the protein system. To equalize the two systems’ lipid inventories, lipid counts in the control tube were matched per leaflet and per species to the protein system’s minimized structure by deleting 394 lipids and cholesterol molecules and the remainder distributed among the phospholipids and PI3P in the same proportion as the protein system’s own clash-based removal. This construction guarantees that the control membrane tube differs from protein-filament tube only in the presence of the filament and its docked PI3P, and not in tube radius, box geometry, or lipid inventory, which is the pairing used for all comparisons between the two systems.

### Molecular dynamics simulation analysis

Visual Molecular Dynamics (VMD)^61^ was used for trajectory visualization. All analyses were performed with MDAnalysis 2.10.0^62^ and plotted with matplotlib, using the solvent-stripped production trajectories (three replicates per system, 15 µs each, 1 ns/frame). Lipids were tracked by their headgroup phosphate bead and the filament by its backbone beads, with all distances evaluated under periodic boundary conditions on the production trajectories directly.

#### Axial crossing rate

Lipid crossings of the filament along the tube axis (*z*) were counted as transition paths^63^ between core regions defined along the periodic axial coordinate (box length *L*, medians 61.5 nm for the protein system and 60.5 nm for the control tube), with positions given in the filament frame. *(i) Filament band (F).* The filament’s axial extent was defined as the 1st–99th percentile of the axial positions of lipid headgroup beads found within 0.6 nm (geodesic surface distance) of the filament, pooled over the three replicates: F = [15.3, 46.2] nm (30.8 nm wide). In the control tube, which has no filament, a band of the same width centered in the box was used, F = [14.9, 45.7] nm. The remainder of the tube is a filament-free gap of 30.7 nm (control, 29.7 nm) that is continuous across the periodic boundary. *(ii) Seam strip (P).* A 2.0 nm strip centered on the midpoint of this gap, which in both systems coincides with the periodic boundary: P = [*L* − 1.0, *L*) ∪ [0, 1.0) nm, i.e. [60.5, 61.5) ∪ [0, 1.0) nm (control, [59.5, 60.5) ∪ [0, 1.0) nm). P marks where a lipid passes around the tube through the gap and the periodic boundary rather than through the band. *(iii) Core regions below (A) and above (B) the band, between P and F*: A = [1.0, 15.3) nm and B = (46.2, 60.5] nm, 14.3 nm each (control, A = [1.0, 14.7) nm and B = (45.7, 59.6] nm, 13.9 nm each). Region boundaries were kept at fixed fractions of the box length, so they follow its fluctuations (∼0.1%). Each lipid was followed along an unwrapped axial coordinate, obtained by wrapping every frame-to-frame displacement of its headgroup bead into [−*L*/2, *L*/2) and summing, so that the periodic boundary itself is never an event. In this coordinate every frame spent in a core region was assigned to that region’s periodic image *n* (A*ₙ* or B*ₙ*). A transition path runs from the last frame a lipid spends in one core region to the first frame it reaches a core region again. A path from A*ₙ* to B*ₙ* (or from B*ₙ* to A*ₙ*) traverses F and was counted as a crossing (upward or downward). A path from B*ₙ* to A*ₙ*h₁ (or from A*ₙ* to B*ₙ*₋₁) traverses P and the gap and was not counted, and a path returning to the region it started from (an entry into F or P that turned back) was not counted either. As a consistency check, every crossing had to contain at least one frame in F, every passage through P none, and no path could change periodic image by more than one region; no violations occurred. Each crossing was attributed to the leaflet in which it occurred, using the per-frame leaflet assignment: a path counted for a leaflet if it started and ended in that leaflet and at most 5% of its leaflet-assigned frames were in the other leaflet; paths during which the lipid switched leaflet were counted separately as leaflet- switching transits. Crossing rates are reported per species and leaflet as the number of crossings divided by the trajectory length (15 µs), as events per µs, mean ± s.d. over the three replicates. The 56 PI3P molecules docked into the filament’s FYVE pockets were excluded from this and every other lipid statistic, since they are considered a protein-bound ligand rather than a leaflet lipid; only the free (bulk) PI3P pool was used. *Filament-frame alignment.* To analyze lipid organization relative to the filament, each protein-system trajectory was aligned to the filament by a translation and a rotation about the tube (*z*) axis only. In each frame, the centroid of all filament beads was superimposed on that of the reference, and the rotation angle about *z* that minimizes the mean-square deviation of the filament beads was obtained in closed form as φ = atan2(H₁₂ − H₂₁, H₁₁ + H₂₂), where H = YᵀWX is the (weighted) cross-covariance matrix of the centered mobile (Y) and reference (X) coordinates. Rotations were determined incrementally between consecutive frames and summed; fitting every frame directly to the reference gave the same angle to within 1.1° over the full 15 µs. The transformation was applied to all atoms. Because the three replicates start from an identical structure, aligning each to its own first frame places all three in a common filament frame. Over 15 µs, the filament rotated about the tube axis by −149°, +72° and −19° in the three replicates. The control tube, which contains no protein, was analyzed without alignment. *Tube diameter.* For each frame, the tube was divided into 32 axial slabs (∼2 nm each), and a circle fit^64^ was applied to the lipid headgroup beads of the inner and outer leaflet in each slab to obtain per-slab leaflet radii and a local tube centerline, which was smoothed over 3 neighboring slabs. Leaflet head radii were obtained from the bimodal radial distribution of headgroup beads about the centerline. Lipids were assigned to leaflets in every frame relative to the local bilayer midplane: in each slab, the inner and outer phosphate surfaces were fitted as functions of the azimuthal angle θ, r(θ) = r₀ + Σₕ₌₁,₂ (aₕ cos hθ + bₕ sin hθ), using phospholipid phosphate beads only, and a lipid was assigned to the inner or outer leaflet when its headgroup bead lay more than 0.3 nm inside or outside the midpoint of the two local surfaces, and was left unassigned otherwise; labels were smoothed with a five-frame running median. Fitting the surfaces as functions of θ allows the boundary to follow the filament-induced, non-circular deformation of the tube cross-section, and placing it at the local midplane keeps it clear of cholesterol’s headgroup bead, which sits ∼0.5 nm below the inner phosphate plane. To compare the dimensions of the two systems directly, the inner- and outer-leaflet radii were converted to diameters (2r) and tracked over the entire production trajectory from t = 0, subsampled every 50 ns and averaged (mean ± s.d.) over the three replicates of each system; the protein system and control tube were overlaid on the same axes for this comparison. Before pooling statistics across the production window, each replicate was checked against five drift criteria evaluated over the analysis window (drift in lumen radius, far-band outer-leaflet lipid count, and undulation amplitude; far-field sampling yield; and fraction of unassigned lipids), to confirm that the tube geometry had reached a stationary state.

#### Filament constriction and per-tetramer RMSD

Filament constriction over time was quantified as the mean radial distance of all filament backbone beads to the local tube centerline.

#### Per-species density maps

Lipid headgroup positions were unrolled onto the tube surface as an arclength (azimuthal) and axial-position pair and binned into a two-dimensional histogram per lipid species over the trajectory’s second half. Each species’ map was normalized by its own mean per-bin occupancy, so that a value of 1 corresponds to the species’ average density on the tube. For the protein filament system, the maps were computed from the filament-aligned trajectories, so that the three replicates are pooled in a common filament frame and the maps report lipid density relative to the filament.

#### Leaflet-resolved lipid enrichment

For each lipid species and leaflet (assigned per frame as above), the pair distribution relative to the filament was computed as a function of geodesic surface distance to the filament’s membrane-proximal footprint (centroids of filament residues with beads within 1 nm of the outer headgroup plane), using the true (unrolled-surface) annulus area at each distance over the trajectory’s final third. Lipids within 5.5 nm of the porin were excluded from this calculation at each frame, so that the porin’s own local perturbation of the membrane did not confound the filament-proximity signal. The 56 PI3P molecules docked into the filament’s FYVE domains were excluded. This was expressed as the enrichment factor ρ(d)/ρ_far_, where ρ(d) is the local number density of the species at surface distance d and ρ_far_ is its density far from the filament (mean over the three outermost distance bins); a value of 1 is the no-enrichment reference, values above 1 indicate local density in excess of the far-field value and values below 1 depletion. Because the leaflets do not sit at a uniform radius around the tube (beneath the filament, inner-leaflet headgroups lie at ∼0.8 of that leaflet’s median radius), each distance shell’s area was evaluated at the median headgroup radius of all lipids of that leaflet within the shell, rather than at the leaflet’s median radius over the whole tube; without this correction, the narrower inner leaflet beneath the filament appears artifactually depleted. Every species normalized to its own far-field density also inherits any change in total lipid density near the filament, which affects all species alike; to isolate species-specific preferences, a relative enrichment was computed as each species’ enrichment factor divided by that of all lipids of the same leaflet pooled, so that a value of 1 indicates that a species follows the local lipid density. Values were computed per replicate and reported as mean ± s.d. over the three replicates for each system.

#### Leaflet composition over time

The mol fraction of each lipid species (cholesterol included) in each leaflet was computed in every frame as the number of lipids of that species assigned to the leaflet divided by the number of all lipids assigned to it, using the per-frame leaflet assignment described above. The 56 docked PI3P were excluded, and cholesterol left unassigned at the bilayer midplane were counted toward neither leaflet. Values were averaged over consecutive 50 ns blocks across the full production run and reported as mean ± s.d. over the three replicates of each system.

#### Lipids removed to place the filament

The lipids deleted during system construction were attributed to individual protomers by following the clash-based removal (any lipid with a bead within 0.4 nm of a filament, docked-PI3P or porin bead; non-periodic distance search). Each deleted lipid was assigned to the leaflet it belonged to in the unperturbed tube, and to a cause: a clash with the filament (assigned to the protomer of the nearest filament bead), a clash with a docked PI3P only (assigned to the protomer whose pocket held that PI3P), or a clash with the porin only (not assigned to a protomer). The five lipids removed afterwards to eliminate residual overlaps, e.g., with other lipids, were assigned to the protomer of the nearest filament bead, and the lipids removed to insert the porin were counted separately (not shown). The footprint of each protomer on the outer leaflet was defined as its projected shadow: every bead of the protomer was projected radially onto the outer headgroup surface of the bare tube (a cylinder of radius r_out_ = 5.6 nm, the median radius of the outer-leaflet phosphate beads), giving a point on the unrolled surface (arc length r_out_·θ, axial position *z*; periodic in both), and the footprint is the union of discs of radius 0.26 nm (half the Martini 3 regular-bead contact distance, 2^1/6^σ with σ = 0.47 nm) around these points. The lipids under the footprint were counted as the outer-leaflet headgroup beads of the tube (before removal) falling inside it. The share of the footprint’s lipids that was removed is reported per protomer and pooled over the 28 protomers (total removed / total under the footprints).

#### Penetration depth of the filament

The depth of each filament bead below the membrane surface was measured in every frame of the three filament-aligned replicates against the local outer lipid-headgroup surface. For each filament bead within 1.5 nm (radially) of the outer phosphate surface, a plane r = a + b·s + c·z was least-squares fitted to the outer-leaflet phospholipid phosphate beads lying 0.8–2.0 nm from the bead on the unrolled surface (s = R·θ, with R the frame’s median outer-leaflet radius; periodic in s and z; docked PI3P and cholesterol excluded), and the depth was taken as (a − r_bead_)/√(1 + b^2^ + c^2^), i.e. the distance along the local surface normal, positive inside the membrane. The inner radius of the annulus excludes the phosphate beads displaced by the bead itself, and fitting locally lets the reference follow deformations of the membrane under each protomer. Beads within 5.5 nm (geodesic) of the porin, where the membrane follows the porin’s rim rather than the tube, were excluded, and frames in which any bead lay more than 2.5 nm below either reference (a failed surface fit; one frame of 45,000) were discarded. For each protomer and frame we recorded the depth of the protomer’s deepest bead and of its deepest backbone bead, and for each residue the depth of its deepest bead and of its backbone bead. The time course of insertion is reported as the per-protomer maxima averaged over the 28 protomers in each frame, which places it on the same per-protomer footing as the per-residue depths. Per-residue depths were pooled over the 28 protomers, all frames and the three replicates and reported as mean ± s.d.; for residues that remain within the analyzed 1.5 nm shell, mean and median agree to within 0.02 nm, and residues whose deepest bead left the shell in more than 5% of frames are not shown. The membrane’s own layers were measured in the same way, every 100 ns, for outer-leaflet phospholipids more than 3 nm (geodesic) from the filament: glycerol beads (GL1/GL2) at 0.35–0.40 nm, the first acyl-chain beads (C1A/C1B) at 0.67–0.72 nm, and the acyl-chain ends (C4A/C4B) at 1.69–1.78 nm below the headgroup surface, with the bilayer midplane at 1.93 nm. *PI3P acyl-chain position and FYVE-domain occupancy.* All outer-leaflet PI3P (the 56 docked and the free molecules) and 150 randomly chosen POPC (reference) were analyzed in the starting structure, every 1 ns for the first 300 ns and every 5 ns thereafter. Each PI3P-binding pocket was defined by the residues within 0.6 nm of the docked PI3P headgroup (inositol C1–C4, PO4 and P3 beads) in at least half of the 28 protomers at t = 0 (FYVE1 pocket: Thr583, Thr587, Trp595, Thr616, Lys617–His619, Arg621, Arg679, Gln686; FYVE2 pocket: Asn565, Ala568, Gln569, Lys704, Trp712, Ser733–His736, Arg738, Arg765). In each frame, a PI3P was classified as pocket-bound if a headgroup bead was within 0.6 nm of a pocket residue (and remained so until it moved beyond 0.8 nm, to avoid counting fluctuations as unbinding), as protein-associated if a headgroup bead was within 0.6 nm of any other filament residue, and as bulk otherwise. FYVE-domain occupancy was the fraction of the 28 FYVE1 (residues 595–656) and 28 FYVE2 (residues 712–772) domains with any PI3P headgroup bead within 0.6 nm, averaged over the full run. The position of the acyl chains was measured relative to the lipid’s own phosphate bead and relative to the local outer headgroup surface (as above, excluding the lipid’s own phosphate). The contact frequency of each filament residue was the fraction of frames, every 5 ns, in which an acyl-chain bead of any PI3P was within 0.6 nm of the residue, averaged over its 28 copies. All quantities are reported as mean ± s.d. over the three replicates.

### Cell culture and stable cell lines

Lysosome-damaging agents used in this study: L-leucyl-L-leucine methyl ester (cat. no. 16008) used at 250 μM, Gly-Phe-β-naphthylamide (cat. no. 14634) used at 250 μM. LysoTracker Deep Red (cat. no. 12492) for live-cell imaging was purchased from Molecular Probes and was used at 75 nM. Janelia Fluor 646 HaloTag ligand (cat no. HT1060) was purchased from Promega and was used at 0.5 uM.

Human U2OS cells (ATCC: HTB-96) were maintained in DMEM (Gibco) medium, supplemented with 10% fetal bovine serum (FBS), 5 U/ml penicillin, and 50 μg/ml streptomycin. Cells were maintained at 37°C supplemented with 5% CO_2_. U2OS-DFCP1-KO cells were previously published ^15^.

Third generation lentiviruses were generated using procedures and plasmids as previously described ^65^. Briefly, tagged fusions of transgenes were generated as Gateway-compatible pENTR constructs using standard molecular biology techniques. From these vectors, lentiviral transfer vectors were generated by Gateway LR recombination into the lentiviral destination vector pLenti-PGK-Blast-DEST (Addgene plasmid #19065, a gift from Eric Campeau and Paul Kaufman). VSV-G pseudotyped lentiviral particles were packaged using a third-generation packaging system (Addgene plasmids #12251, #12253, and #12259, a gift from Didier Trono). Cells were then transduced with low virus titers and stable expressing populations were generated, either by antibiotic resistance and flow cytometry. The weak PGK promoter was used for transgene expression in order to achieve low expression levels.

### Plasmid constructs

VPS13C-Halo was a gift from Pietro De Camilli (Addgene plasmid #232864 ^10^), and mCherry-DFCP1 was described before ^15^. Using standard molecular cloning techniques we generated the following DFCP1 mutants: membrane-bound tetramer interface mutant: R458D/R536D/I539D; membrane embedding mutant: W504D/W524D/F525D/W585D; PI3P binding mutant: H618D/R621E/H735D/R738E.

### Live-cell microscopy

Cells seeded in MatTek 35-mm petri dish, 20-mm Microwell No. 1.5 coverglass or on Ibidi m-slide 4 well ibiTreat chambers were imaged on a Nikon Eclipse Ti2-E inverted microscope equipped with a CrestOptics X-Light V3 spinning disk confocal module, a dual camera imaging setup (back-illuminated sCMOS cameras, Photometrics Kinetix) and a CFI Plan Apo 100X Oil *NA 1.45* lens. For temperature control during live observation, the microscope stage was kept at 37°C by a temperature-controlled incubation chamber. Cells were imaged in live-cell imaging buffer (Invitrogen, supplemented with FBS, penicillin/streptomycin, HEPES and Aln-Gln) with 20 mM glucose, at 37°C and 5% CO_2_.

### Image analysis and post-processing

For LysoTracker recovery analysis, a fully automated analysis using NisElements software, detecting LysoTracker positive bright puncta per timepoint, from live cell imaging movies was applied ^12^. Intensity-based thresholding was used for segmentation of cell area. Identical settings for all conditions within one experiment. The analysis of lysosome damage response of VPS13C in cells expressing WT or mutant DFCP1 was automated by using the NisElements software, detecting bright puncta of VPS13C-Halo in single cells per timepoint. To account for the variation in expression level of VPS13-C-Halo in transiently transfected cells, the mean intensity of VPS13C-Halo was measured in the whole cell, and the cells were then assigned to a dedicated analysis, set for low, medium or high expression. Identical selection criteria were used for WT and mutant in all experiments.

### Statistics and reproducibility

The number of individual experiments and the number of cells analyzed are indicated in the figure legends. Graphs were created using GraphPad Prism 10. No statistical methods were used to predetermine sample sizes. The statistical tests performed for each experiment are specified in the figure legends. P-values are indicated for each experiment. Data collection and analysis were not performed blind to the conditions of the experiments and no randomization was applied. Data were excluded only for technical reasons, including loss of focus, excessive photobleaching, or phototoxicity during live-cell imaging experiments. To ensure the reproducibility of the data presented, experiments were performed by different researchers.

## Acknowledgements

We thank R. Hooy and K. Sharma of Cal Cryo for cryo-EM and cryo-ET support. This research was funded by Aligning Science Across Parkinson’s (ASAP-000350) through the Michael J. Fox Foundation for Parkinson’s Research (to G. H. and J.H.H.), with seed funding from the Peder Sather Foundation (H. S. and J. H.). K.P.-R. and G.H. thank the Max Planck Society for support. C. R. was supported by a grant from the Norwegian Cancer Society (project number 246670). M. R. was supported by a Young Research Talents Grant from the Research Council of Norway (grant number 345137) and Norwegian Cancer Society (grant number 322550). H. S. was supported by grants from the Norwegian Cancer Society (project number 182698), the South-Eastern Norway Regional Health Authority (project number 2016087), the Research Council of Norway (project number 302994), the European Research Council (Advanced Grant number 788954), and the Miller Foundation of UC Berkeley.

## Author Contributions

K.T. and Z. C. purified proteins. Z. C. carried out liposome reconstitutions and single-particle cryo-EM. E. H. carried out cryo-ET. K. P.-R. carried out molecular dynamics simulations. A. K. performed DFCP1 mutagenesis and molecular cloning and generated the stable cell lines. M. R. and S. P. carried out the light microscopy and lysosomal repair assays. C. R. performed the image analysis and statistics. The project was initially conceived by J. H. and H. S., and supervised by J. H., H. S., G. H., and Z. C. Z. C. and J. H. wrote the first draft of the manuscript, which was checked and edited by all authors.

## Data availability

The cryo-EM and cryo-ET maps have been deposited in the Electron Microscopy Data Bank (EMDB) under the following accession codes: five-tetramer higher-order assembly map (EMD-79488), focused map of the tetrameric ATPase domain with ERTD (EMD-79378), focused map of the monomeric FYVE domain with ERTD (EMD-79380), composite map (EMD-79404), and the tetrameric map with C2 symmetry (EMD-79381). The atomic coordinates for the DFCP1 tetramer have been deposited in the Protein Data Bank (PDB) under accession code 39FP. The MD data have been deposited in a Zenodo repository that will become public upon acceptance (with access by code to editors and reviewers). All other data are available from the corresponding authors upon reasonable request.

## Extended Data Figures

**Extended Data Figure 1.**
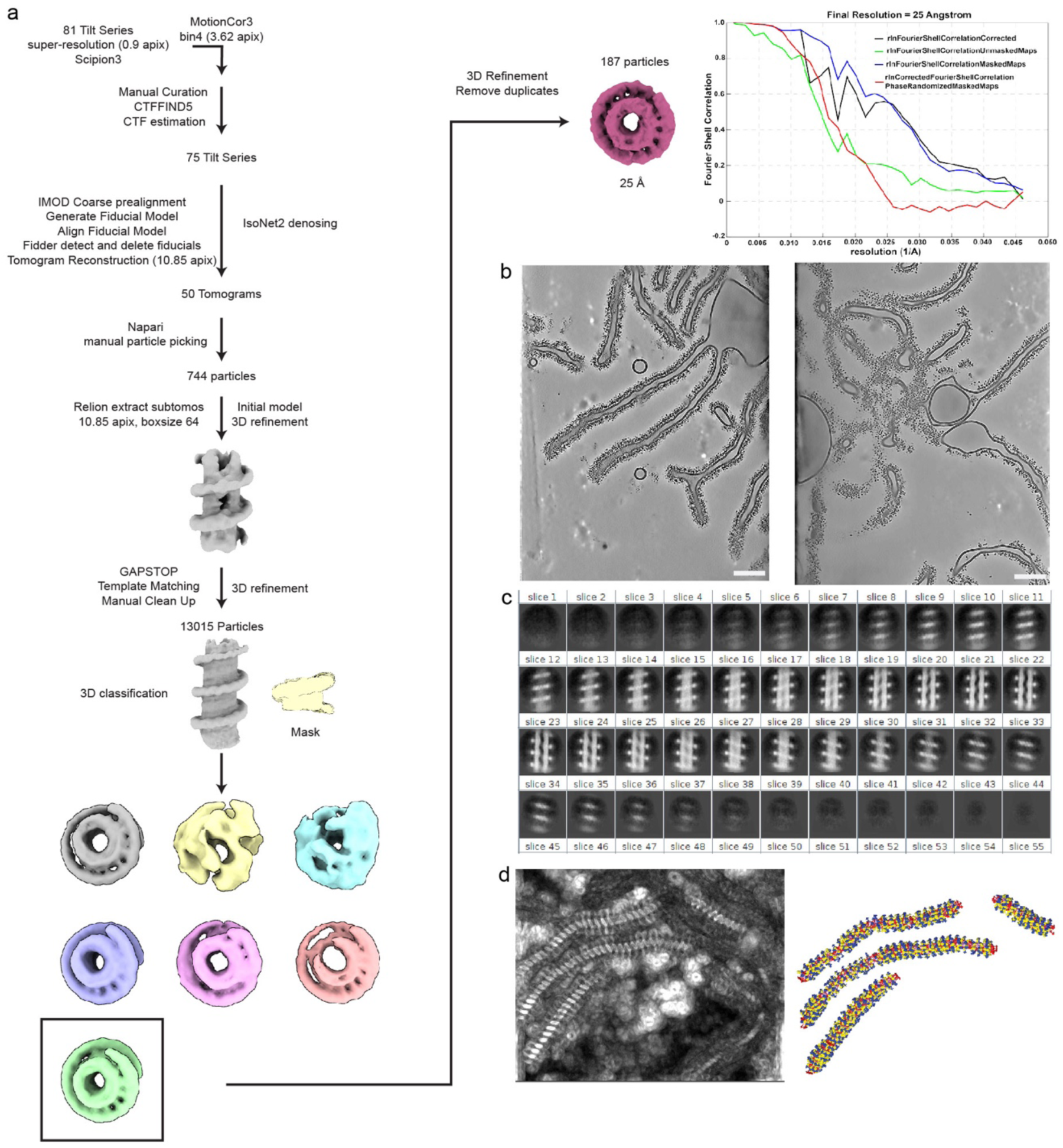
Cryo-ET data processing and structural refinement workflow. **a**, Schematic of the cryo-ET workflow, from tilt-series alignment in IMOD to sub-tomogram averaging. **b**, Representative tomograms showing DFCP1-induced tubulation of spherical liposomes (left) and the fuzzy, non-fully constricted assembly (right). Scale bar: 100 nm. **c**, 2D slices of the density map after 3D refinement, displaying the arrangement of DFCP1 around the tubule along the Y-Axis. (d) Template matching (TM) score map showing the lattice assembly along the tubules (left) and the extracted coordinates with Euler angles following the mechanical twist (right).

**Extended Data Figure 2.**
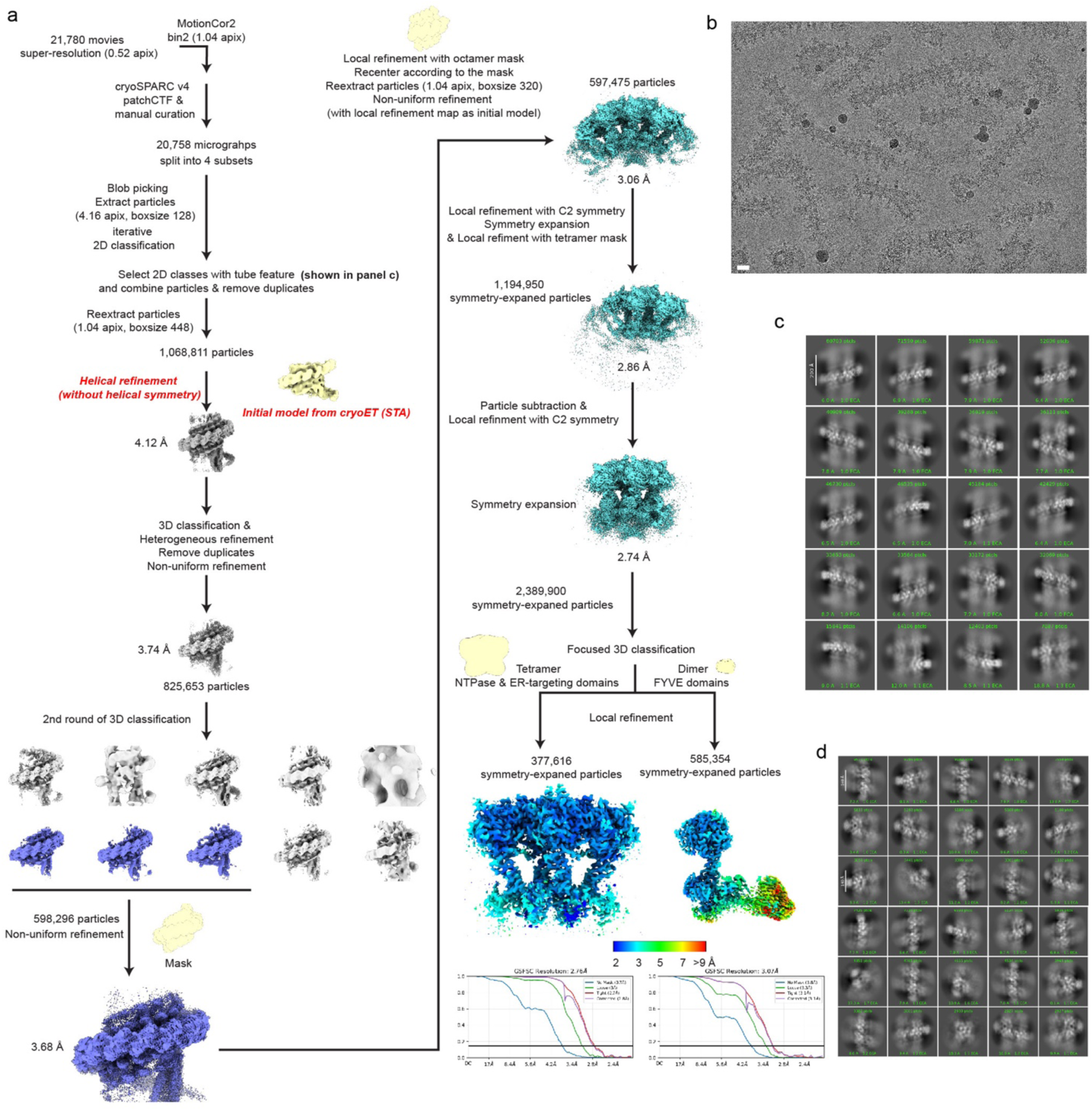
Single-particle cryo-EM data processing workflow. **a**, Schematic of the single-particle cryo-EM workflow. **b**, A representative image of DFCP1 with liposomes. **c**, Representative 2D class averages showing heterogenous tube structure. **d**, Representative 2D class averages showing high-resolution features of the final particle stack focusing on the octamer assembly with a box size of 320.

**Extended Data Figure 3.**
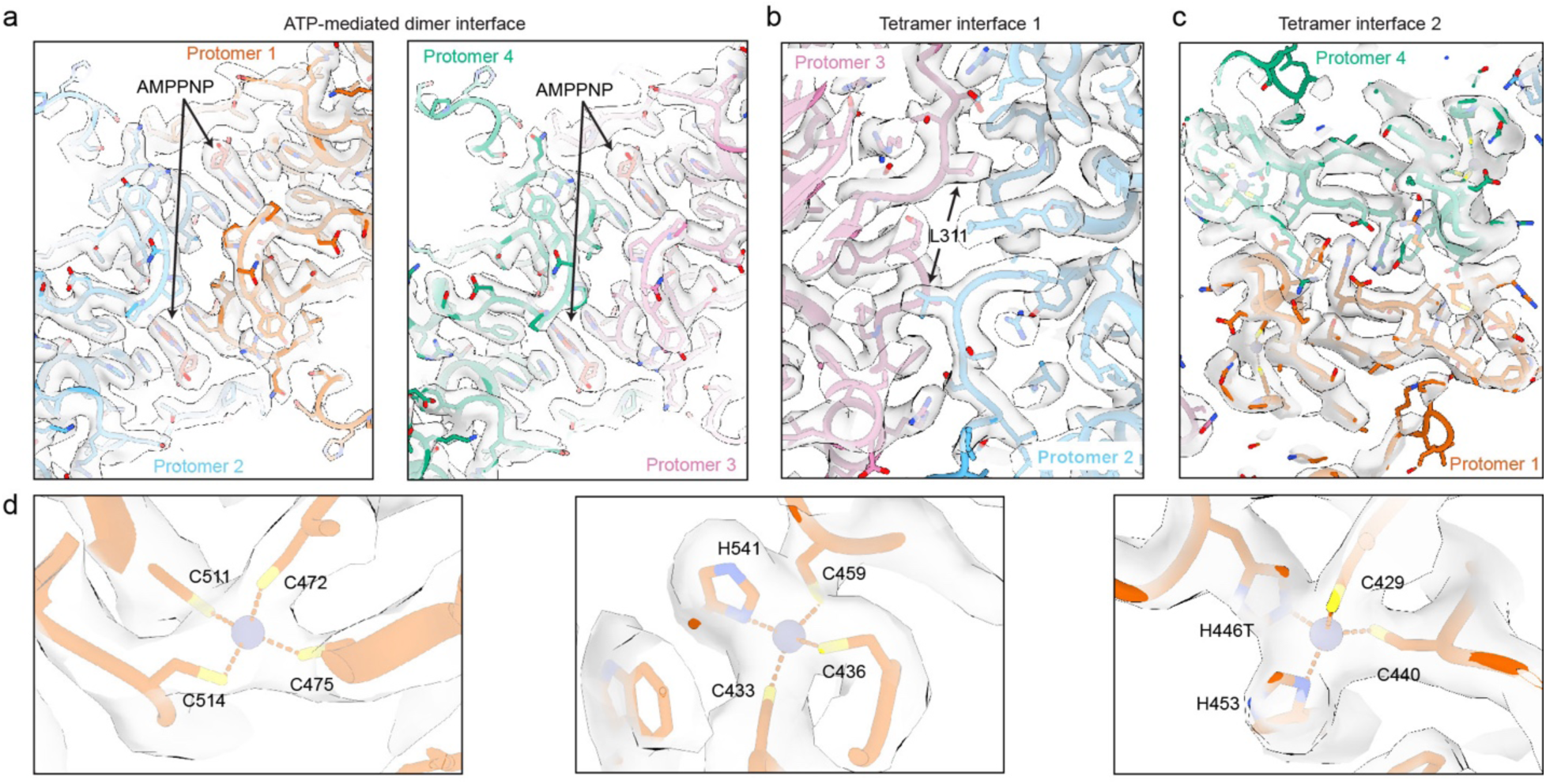
Cryo-EM density examples and atomic model fitting. Representative density for the nucleotide-mediated dimerization interface (**a**), tetramer interface (**b**,**c**), and three structural zinc-binding sites (**d**).

**Extended Data Figure 4.**
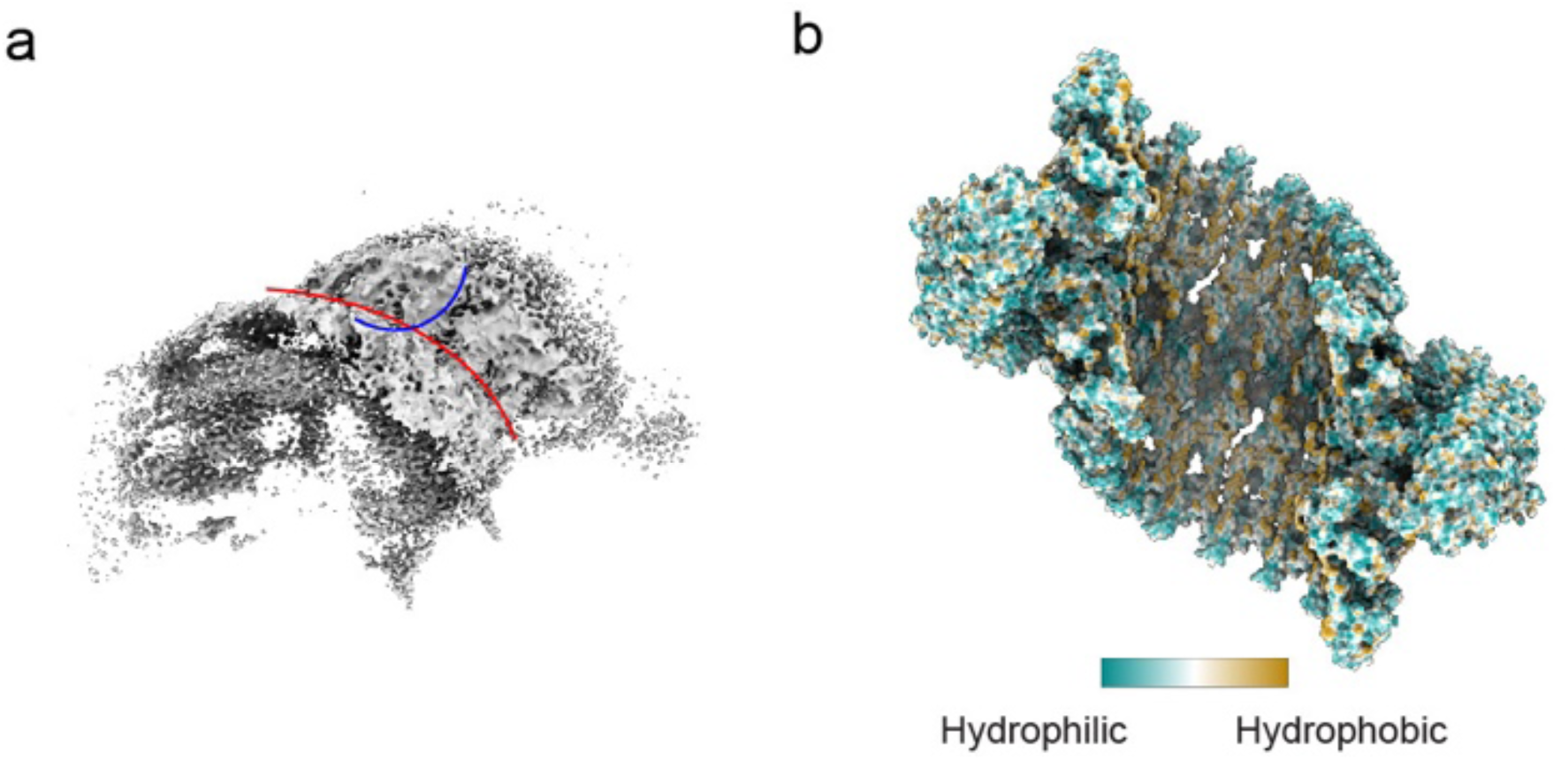
Cryo-EM analysis of membrane geometry. **a**, Segmented cryo-EM map of the membrane from the DFCP1 oligomer map. The geometry highlights the distinct local curvatures of the membrane bilayer at the protein docking site, showing a saddle-like shape. **b**, Hydrophobicity map of the higher-order assembly consisting of five adjacent tetramers. The surface is colored by hydrophobicity, illustrating the extensive hydrophobic footprint that anchors the filamentous array into the membrane.

**Extended Data Figure 5.**
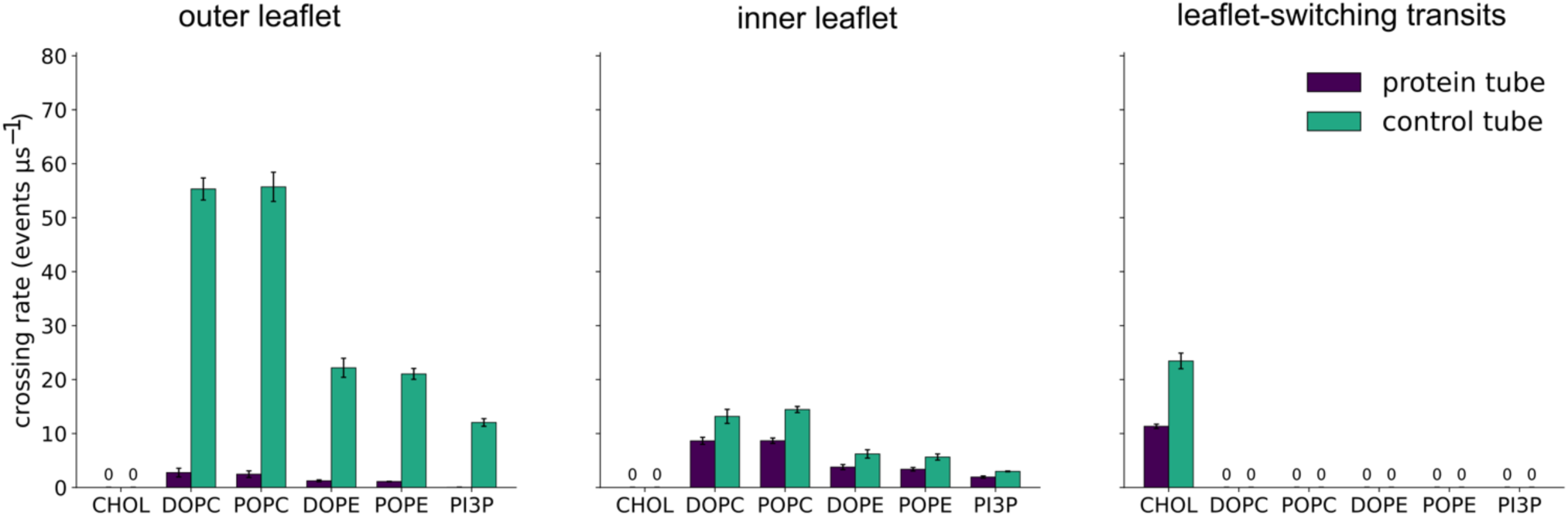
Leaflet-resolved axial crossing rate. The same axial crossing rate as Fig. 3c, broken down by which leaflet the lipid travelled in: within the outer leaflet (left), within the inner leaflet (middle), and for transits during which the lipid switched leaflet on the way (right), for the filament (dark purple) and in the control tube, for a band of the same axial width placed at the control tube’s center (green), reported as crossing events per µs of trajectory (mean ± s.d., n = 3 replicates per system). Cholesterol crosses only while switching leaflets, owing to its fast flip-flop, so its rate is ∼0 in the outer- and inner-leaflet panels; zero rates are marked “0”.

**Extended Data Figure 6.**
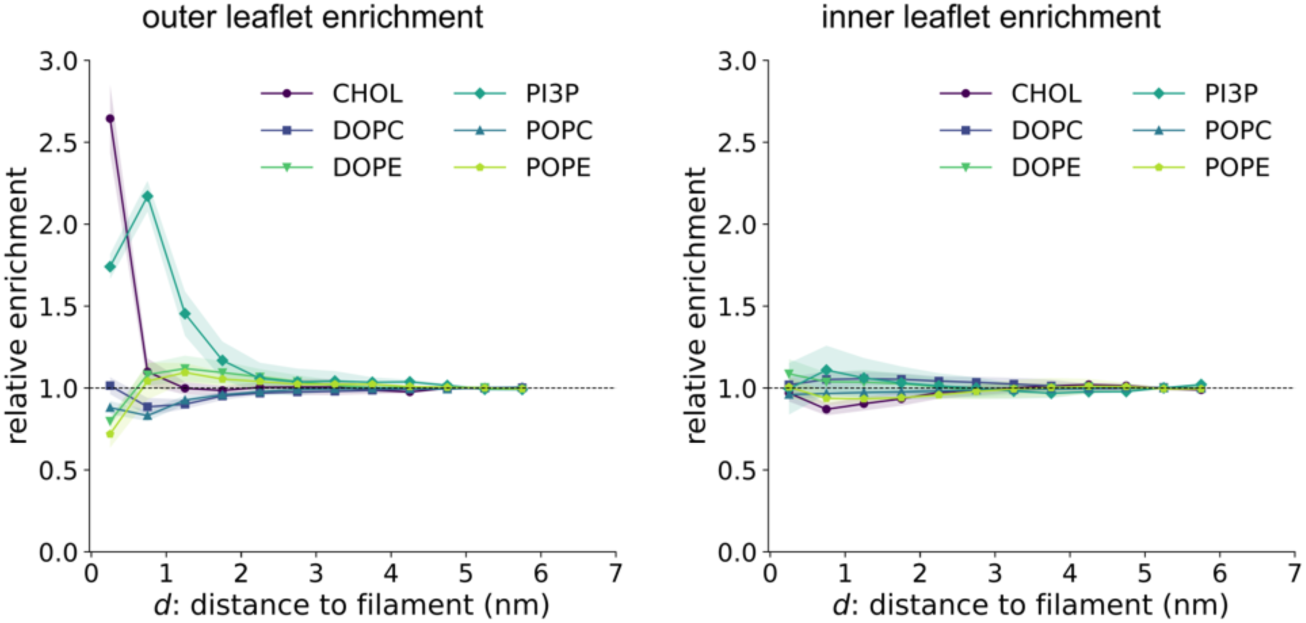
Leaflet-resolved lipid enrichment around the filament. Relative enrichment of each lipid species as a function of geodesic surface distance d to the filament, in the outer (left) and inner (right) leaflet, for the filament system. Relative enrichment factor ρ(d)/ρfar, with shell areas evaluated at the local leaflet radius, divided by that of all lipids of the same leaflet pooled (Methods); it removes the change in total lipid density near the filament that all species share and isolates species-specific preferences, a value of 1 meaning that the species follows the local lipid density. Left, outer leaflet; right, inner leaflet. Each lipid species is shown in its own color (viridis: cholesterol, dark purple; DOPC, indigo; POPC, blue; PI3P, teal; DOPE, green; POPE, yellow-green; markers: cholesterol, circle; DOPC, square; POPC, upward triangle; PI3P, diamond; DOPE, downward triangle; POPE, pentagon), computed over the trajectory’s final third and reported as mean ± s.d. over 3 replicates; the dashed horizontal line marks the no-enrichment reference, ρ(d)/ρfar = 1. Values above 1 indicate recruitment toward the filament and values below 1 depletion.

**Extended Data Figure 7.**
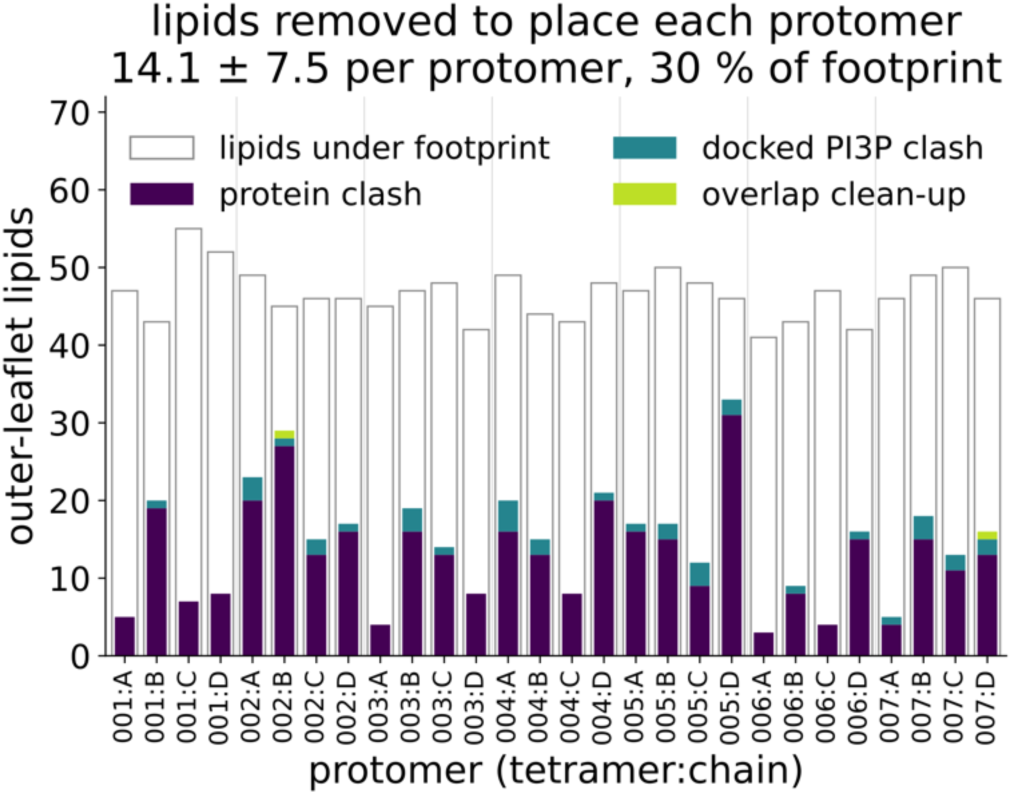
Outer-leaflet lipids removed to place each protomer. Number of outer-leaflet lipids deleted during system construction to accommodate each of the 28 protomers of the filament (protomers labelled tetramer:chain, 001:A–007:D; grey vertical lines separate tetramers). Colored bars stack the lipids removed because they clashed with the protomer itself (protein clash, dark purple), with the PI3P docked in the protomer’s His/Arg FYVE domains pockets but not the protein (docked PI3P clash, teal), or because they still overlapped any other component of the system (e.g., other lipids) before energy minimization (overlap clean-up, yellow-green). Open bars give the number of outer-leaflet lipids that lay under the protomer’s footprint in the tube before removal, the footprint being the protomer’s shadow projected radially onto the outer headgroup surface (Methods). Lipids removed to insert the porin, or for clashing with the porin only, are not attributed to any protomer and are not shown. The title gives the mean ± s.d. over the 28 protomers and the pooled share of the footprint’s lipids that was removed.

## Supplementary Information

### Extended Analysis of MD Simulations

We built coarse-grained Martini 3 models of two systems, the DFCP1 filament wrapped around a membrane nanotube and a protein-free tube of matched radius, box geometry and lipid composition. The filament comprised 28 protomers arranged as seven stacked tetramers around the tube axis. Both membrane tubes were built starting from a bilayer of DOPC, POPC, DOPE, POPE, 10% PI3P and cholesterol. For the protein system, 56 PI3P molecules were docked into the paired histidine and arginine pockets of the FYVE domains, two per protomer, before production. Each system was run for 15 µs in three independent replicates.

### Lipids displaced by the filament

The deep insertion of the ER-targeting domain displaces a large number of outer-leaflet lipids. Attributing the clash-based removals to individual protomers gave an average of 14 ± 8 outer-leaflet lipids removed per DFCP1 monomer, about 30% of the lipids lying under each monomer’s footprint (Extended Data Fig. 7). Rapid cholesterol flip-flop between the lipids compensates for remaining packing issues that could have resulted from incomplete or excessive lipid removal from the outer leaflet. However, the lipid mole fraction in inner and outer leaflets closely matched that of the protein-free tube (Supplementary Figure 3) where only the lipids clashing with the solvent-exchange porin had been removed. Note that the lipid inventories of the two systems were matched per leaflet and per species so that the control tube differs from the filament tube only in the presence of the filament and its docked PI3P.

### A leaflet-resolved diffusion barrier

Resolving the crossing rates by leaflet shows that the barrier acts primarily on the outer leaflet in direct contact with the filament. PI3P crossed within the outer leaflet only once in 45 µs of combined sampling, against 12.0 ± 0.7 crossings per µs in the control system, so PI3P mobility is arrested specifically by the filament and not by the tube geometry (Supplementary Fig. 2 and Supplementary Movie 1). The bulk phospholipids showed the same asymmetry, with a far stronger block in the outer leaflet than in the inner one (for example 11.4 ± 0.6 against 68.5 ± 3.3 crossings per µs for DOPC; Fig. 3c, Extended Data Fig. 5). Cholesterol was the exception. Every cholesterol crossing switched leaflet, consistent with its fast flip-flop, allowing cholesterol to bypass the fence by flipping to the inner leaflet. Even with this exclusive route past the filament, cholesterol crossings ran at 48% of the control rate.

### Lipid redistribution around the filament

Per-species density maps show a distinct depletion and enrichment pattern confined to the filament’s footprint in the protein system that is entirely absent on the control tube (Supplementary Fig. 1). The apparent increase in density in some regions of the inner leaflet reflects the local protein-induced deformation of the tube rather than true enrichment. Leaflet-resolved profiles sharpen this picture. Free PI3P was recruited immediately beneath the filament in the outer leaflet, in excess of the PI3P lipids bound to FYVE domains, with an enrichment factor of 2.2 ± 0.1 at 0.5 to 1 nm from the filament, and showed no comparable recruitment in the inner leaflet (Extended Data Fig. 6). Cholesterol showed a sharp, highly localized enrichment directly beneath the filament in the outer leaflet, 2.6 ± 0.3-fold within 0.5 nm, returning to the local density by 1 nm, together with a mild depletion in the inner leaflet. These profiles indicate that the filament captures the lipid it directly binds rather than passively perturbing the whole bilayer. Leaflet composition was otherwise stable over the full trajectory and close to that of the control tube (Supplementary Fig. 3).

### Membrane penetration

The filament penetrates the outer leaflet without reaching the bilayer midplane (Supplementary Fig. 4). In each protomer the deepest bead sits 0.98 nm below the headgroup surface, past the glycerol region and into the acyl chains, and this depth is constant over the 15 µs of MD simulation. The insertion is carried mainly by aromatic residues. Trp524 is the deepest at 0.93 ± 0.18 nm, followed by Trp504, Phe525 and Trp496, together with hydrophobic residues of the helix linking the two FYVE domains, including Val692 and Val685 (Supplementary Fig. 4b). All 28 protomers insert to a similar depth, and no part of the filament reaches the midplane at 1.93 nm.

### PI3P in the FYVE pockets

The PI3P lipids initially docked into the FYVE domains generally stay in the pockets and remain inserted in the bilayer throughout the runs, with occasional rebinding of PI3P to vacated FYVE domains from the bilayer (Supplementary Fig. 5). Averaged over the trajectories, a PI3P headgroup contacts 91 ± 4% of the FYVE1 domains and 65 ± 3% of the FYVE2 domains, almost entirely through the simulations (Supplementary Fig. 5a,b). The acyl chains of pocket-bound PI3P stay extended into the bilayer, ending 1.4 to 1.6 nm below the lipid’s own phosphate, close to bulk PI3P, but the whole lipid is raised relative to the surrounding membrane, so that its glycerol sits level with the neighboring phosphates and its chain ends lie about 0.45 nm less deeply than those of bulk PI3P. The filament therefore lifts the PI3P it binds without pulling its acyl chains out of the bilayer. Those chains contact mainly the hydrophobic residues of the membrane-embedded inter-FYVE helix, with Ile696 and Leu689 touching a PI3P chain in 86% of frames and Leu700 and Val692 in 74 to 76% (Supplementary Fig. 5e).

### Supplementary Figures

**Supplementary Figure 1.**
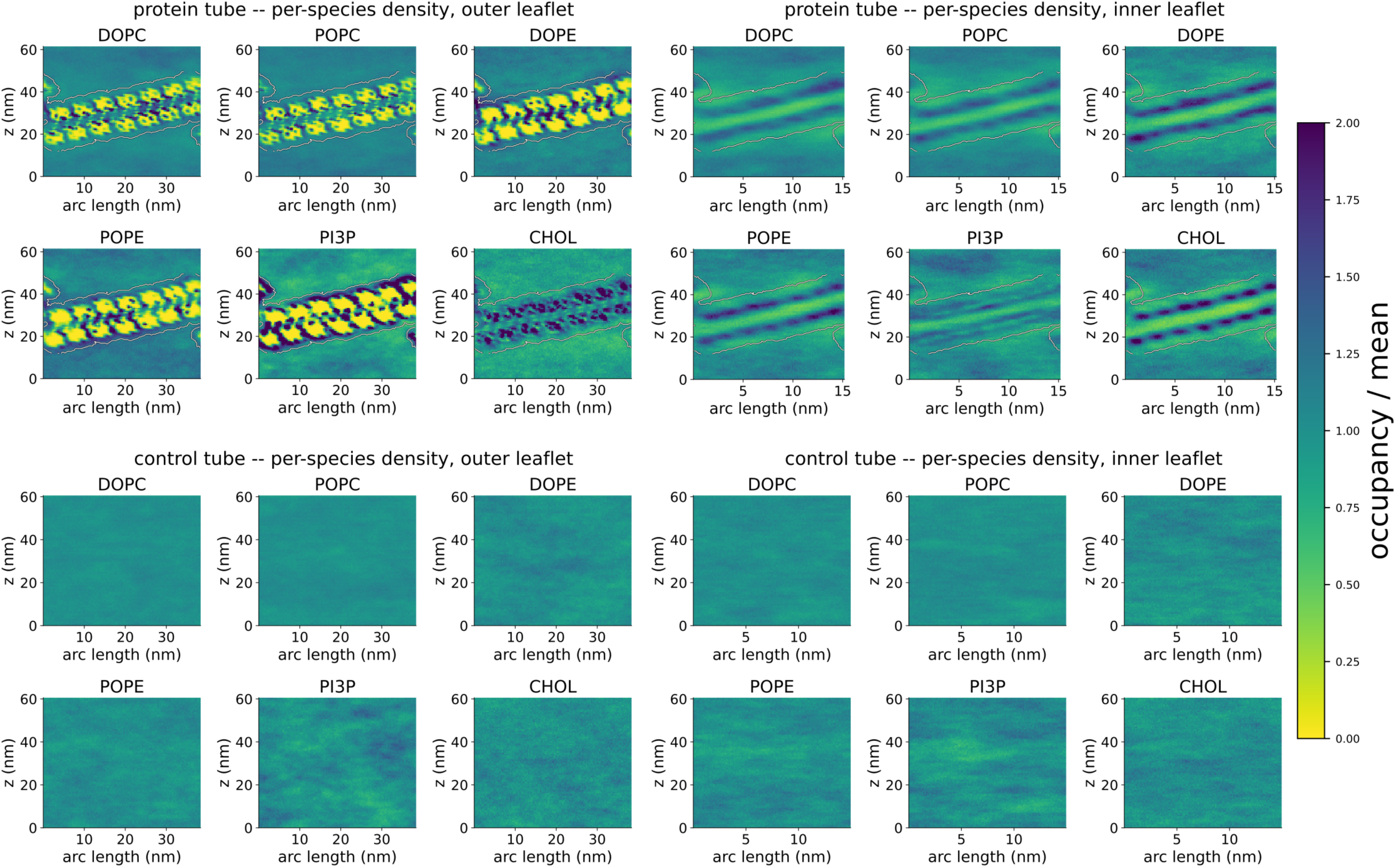
Per-species lipid density on the tube surface. Head-bead occupancy of each lipid species, projected onto the unrolled tube surface (arc length along the circumference vs. axial position, z), pooled over the last half of the production trajectory and all three replicates of each system. Top row, protein filament system; bottom row, protein-free control tube; left column, outer leaflet; right column, inner leaflet. Each species’ panel is normalized by its own mean occupancy, so that a value of 1 corresponds to that species’ average density on the tube; the diverging scale is fixed to [0, 2] across every panel (yellow, depleted; dark purple, enriched). The white outline in the protein system panels marks the filament’s footprint (lipids ever found within 0.6 nm of the filament). For the protein system, the maps were computed from trajectories aligned to the filament (translation and rotation about the tube axis only), so that the footprint and density maps are shown in the filament’s frame of reference rather than the simulation’s lab frame. Lipids are assigned to leaflets in every frame relative to the local bilayer midplane (Methods).

**Supplementary Figure 2.**
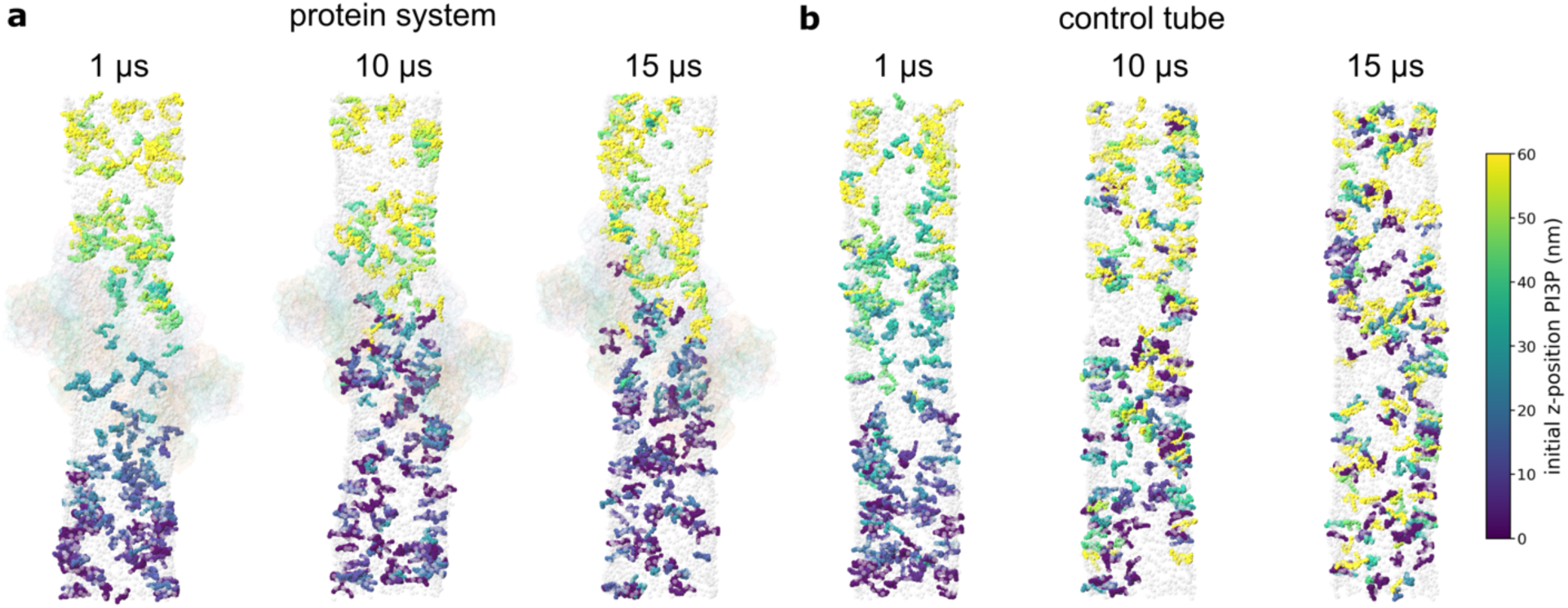
Axial mixing of outer-leaflet PI3P on the tube. **a**-**b**, Snapshots of the filament system (**a**) and the protein-free control tube (**b**) after 1 µs (left), 10 µs (center) and 15 µs (right) of production, viewed from the side with the tube axis vertical (one replicate per system; the full trajectories are shown in Supplementary Movie 1). Each outer-leaflet PI3P lipid is colored by its axial position (z) in the starting structure (viridis, 0 nm dark purple to 60 nm yellow), so that colors that remain sorted along the tube axis indicate limited axial mixing, whereas interspersed colors indicate lipids that have travelled along the tube. Because the simulation box is periodic along the tube axis, lipids leaving through one end of the box and re-entering at the other take the color of the end at which they re-entered, as if they had started there. Lipids are assigned to leaflets in every frame relative to the local bilayer midplane (Methods); inner-leaflet lipids are not shown. The phosphate beads of all other outer-leaflet phospholipids are shown in grey, including those of the PI3P docked in the filament’s FYVE-domain pockets, which are not colored; cholesterol is not shown. The filament is rendered nearly transparent, with its protomer chains colored as in Fig. 3a. For the protein system, the trajectory is aligned to the filament (translation and rotation about the tube axis only), as in Supplementary Fig. 1.

**Supplementary Figure 3.**
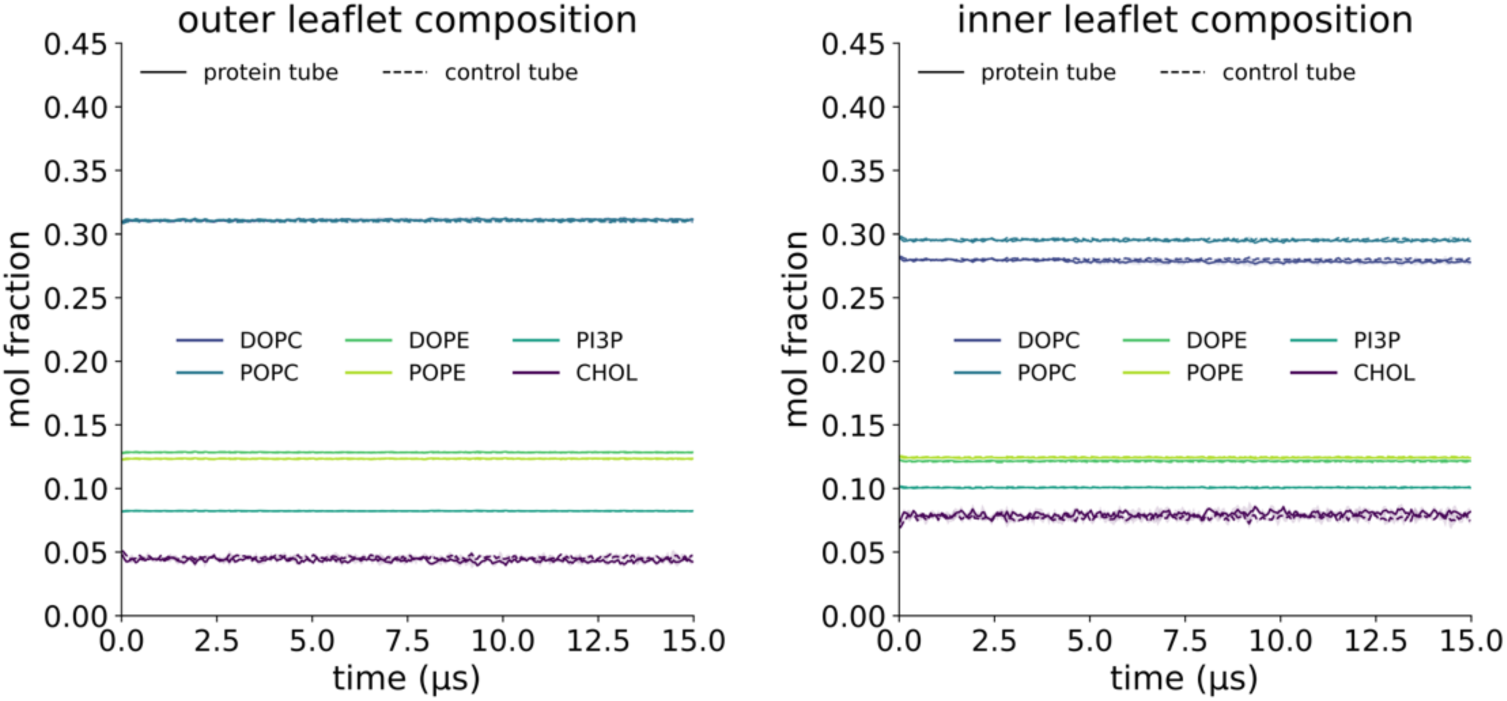
Leaflet lipid composition over time. Mol fraction of each lipid species in the outer (left) and inner (right) leaflet of the tube as a function of time, for the filament system (solid lines) and the protein-free control tube (dashed lines). Each lipid is assigned to a leaflet in every frame relative to the local bilayer midplane (Methods), and the mol fraction is the number of lipids of a species in the leaflet divided by all lipids assigned to that leaflet, excluding the 56 PI3P docked in the filament’s pockets. Values are averaged over consecutive 50 ns blocks; lines show the mean and shaded bands the s.d. over 3 replicates per system. Each lipid species is shown in the same color as in Extended Data Fig. 6 (viridis: cholesterol, dark purple; DOPC, indigo; POPC, blue; PI3P, teal; DOPE, green; POPE, yellow-green).

**Supplementary Figure 4.**
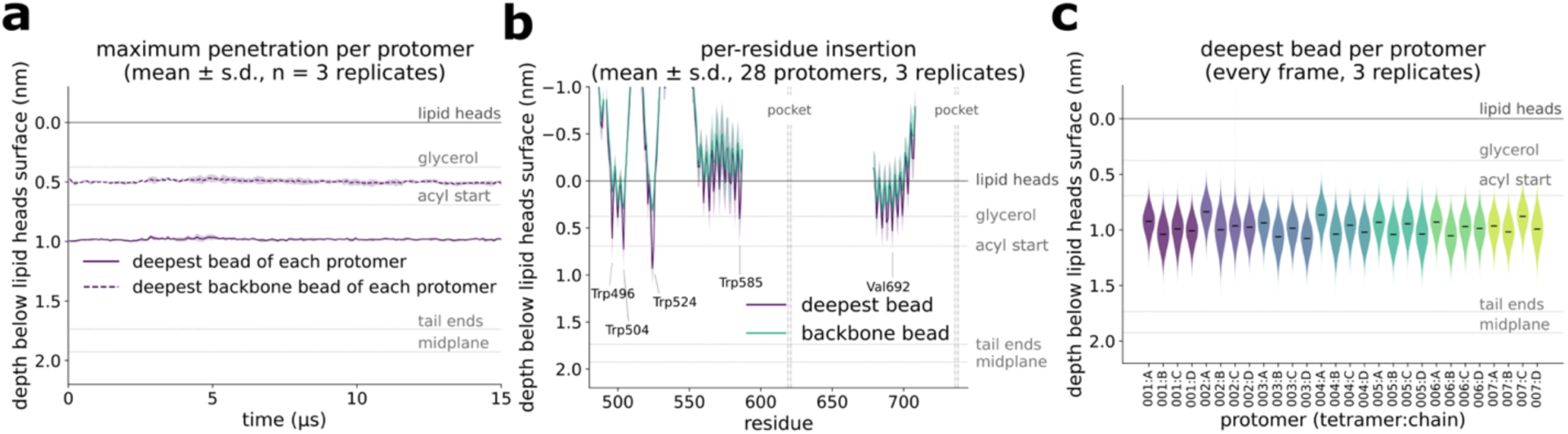
Penetration of the filament into the membrane. Depth below the local outer lipid-headgroup (phosphate) surface, measured along the local surface normal and positive inside the membrane (Methods); in every panel the depth axis grows downward, and the dotted horizontal lines mark the membrane’s own layers measured in the same way far from the filament: glycerol, first acyl-chain bead (acyl start), acyl-chain ends and bilayer midplane; the solid line at 0 is the lipid-headgroup surface. **a**, Depth of each protomer’s deepest bead (solid) and deepest backbone bead (dashed), averaged over the 28 protomers in each frame and over 100 ns blocks, as a function of time; lines show the mean and shaded bands the s.d. over n = 3 replicates. **b**, Depth of each residue’s deepest bead (dark purple) and of its backbone bead (green) along the sequence, for the membrane-contacting region (residues 480–745), pooled over the 28 protomers, all frames and the 3 replicates (line, mean; band, ± s.d.). Residues that do not stay within 1.5 nm of the headgroup surface in at least 95% of frames are not shown; the most deeply inserted residues are labelled, and dash-dotted lines mark the histidine and arginine of the two PI3P-binding pockets. **c**, Distribution of the depth of the deepest bead of each protomer over every frame of the 3 replicates (violins, colored by tetramer; horizontal bars, medians).

**Supplementary Figure 5.**
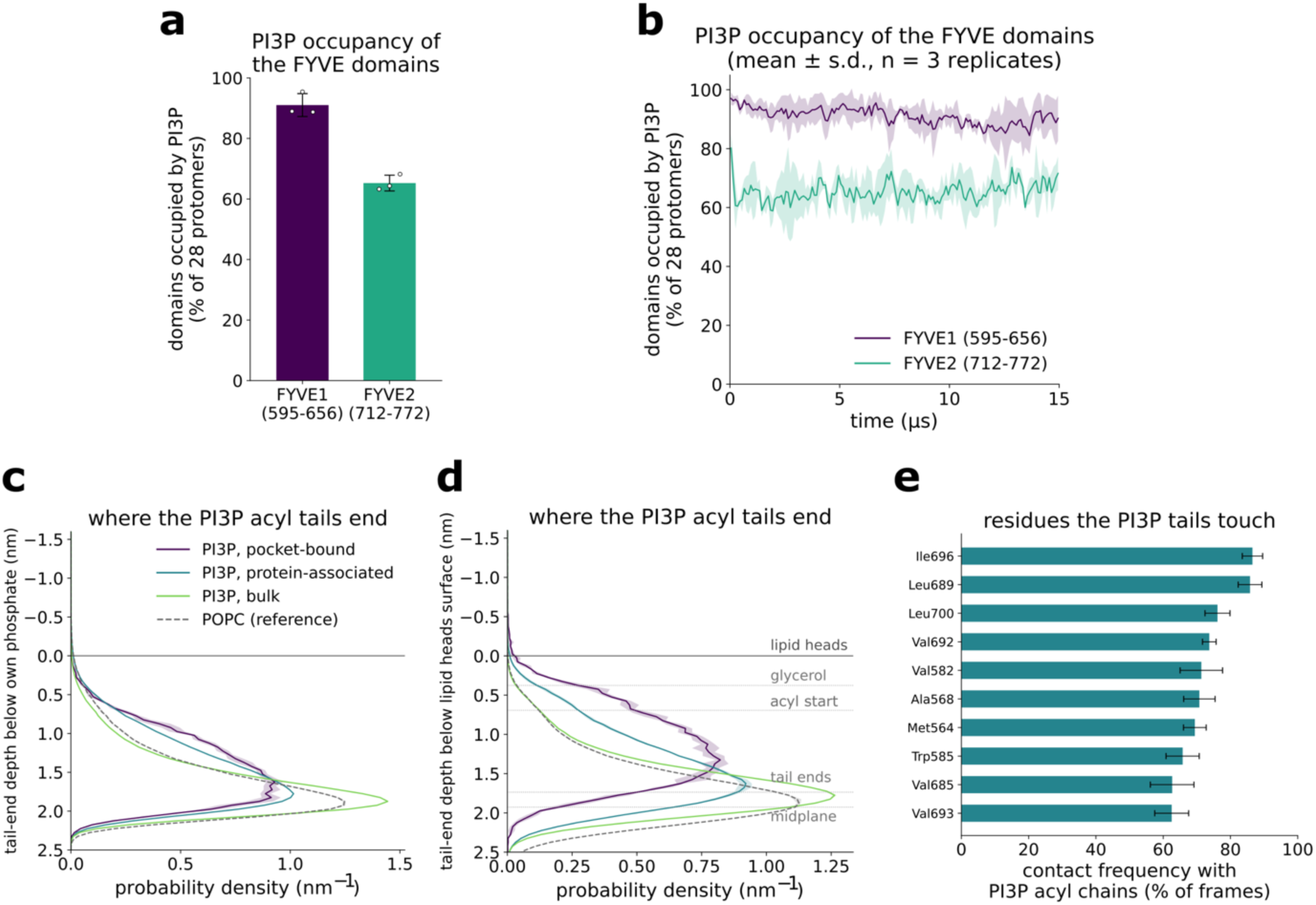
PI3P occupancy of the FYVE domains and acyl-chain position. Each outer-leaflet PI3P was classified in every frame as pocket-bound (headgroup within 0.6 nm of a residue lining one of the 56 PI3P-binding pockets, defined from the docked starting structure; released beyond 0.8 nm), protein-associated (headgroup within 0.6 nm of the filament but not pocket-bound) or bulk (Methods), for panels b and c; POPC of the same leaflet is shown as the bilayer reference. **a**, PI3P occupancy of the two FYVE domains of each protomer, time-averaged over the full 15 µs production run: percentage of the 28 FYVE1 (residues 595–656, dark purple) and 28 FYVE2 (residues 712–772, green) domains with a headgroup bead of any PI3P, docked or free, within 0.6 nm of any domain bead (Methods). Bars and error bars show the mean and s.d. over n = 3 replicates; open circles are the individual replicates. **b**, PI3P occupancy as a function of time for FYVE1 (dark purple) and FYVE2 (green), averaged over 100 ns blocks; lines show the mean and shaded bands the s.d. over n = 3 replicates. **c**, Distribution of the depth of the acyl-chain end beads (C4A and C4B) below the lipid’s own phosphate bead, for pocket-bound (dark purple), protein-associated (teal) and bulk (green) PI3P and for POPC (dashed grey). **d**, As in **b**, with depth measured below the local outer lipid-headgroup surface (Methods); dotted lines mark the membrane layers as in Supplementary Fig. 4. In **c** and **d**, the first 300 ns (every frame) and the remainder of the run (every fifth frame) are pooled; lines show the mean and shaded bands the s.d. over n = 3 replicates. **e**, The 10 filament residues most frequently contacted by the acyl chains of PI3P: contact frequency, the percentage of frames (every 5 ns over the full run) in which at least one PI3P acyl-chain bead, docked or free PI3P, lies within 0.6 nm of the residue, averaged over its 28 copies (one per protomer); bars and error bars show the mean and s.d. over n = 3 replicates.

**Supplementary Movie 1. Axial mixing of outer-leaflet PI3P over 15 µs.** Trajectories of the filament system (left, “protein system”) and the protein-free control tube (right, “control tube”) from the starting structure to 15 µs of production (one replicate per system), rendered as in Supplementary Fig. 2: outer-leaflet PI3P colored by their initial axial position (color bar, 0–60 nm), recolored as described there when they cross the periodic boundary along the tube axis, the phosphate beads of the other outer-leaflet phospholipids in grey, and the filament colored by protomer chain. The movie opens on the starting structure with the filament shown opaque and PI3P hidden; the filament then fades to near-transparency as the PI3P appear, the 15 µs of trajectory play over 10 s (elapsed simulation time at bottom left), and the PI3P finally fade out as the filament returns to full opacity. For the protein system, the trajectory is aligned to the filament (translation and rotation about the tube axis only), as in Supplementary Fig. 1.

## References

1 Ballabio, A. & Bonifacino, J. S. Lysosomes as dynamic regulators of cell and organismal homeostasis. Nat Rev Mol Cell Biol 21, 101–118 (2020). 10.1038/s41580-019-0185-4

2 Shin, H. R. & Zoncu, R. The Lysosome at the Intersection of Cellular Growth and Destruction. Dev Cell 54, 226–238 (2020). 10.1016/j.devcel.2020.06.010

3 Yang, H. & Tan, J. X. Lysosomal quality control: molecular mechanisms and therapeutic implications. Trends Cell Biol 33, 749–764 (2023). 10.1016/j.tcb.2023.01.001

4 Radulovic, M., Yang, C. & Stenmark, H. Lysosomal membrane homeostasis and its importance in physiology and disease. Nat Rev Mol Cell Biol 27, 71–87 (2026). 10.1038/s41580-025-00873-w

5 Skowyra, M. L., Schlesinger, P. H., Naismith, T. V. & Hanson, P. I. Triggered recruitment of ESCRT machinery promotes endolysosomal repair. Science 360 (2018). 10.1126/science.aar5078

6 Radulovic, M. et al. ESCRT-mediated lysosome repair precedes lysophagy and promotes cell survival. EMBO J 37 (2018). 10.15252/embj.201899753

7 Durgan, J. & Florey, O. Many roads lead to CASM: Diverse stimuli of noncanonical autophagy share a unifying molecular mechanism. Sci Adv 8, eabo1274 (2022). 10.1126/sciadv.abo1274

8 Goul, C. S. et al. LASER couples damage sensing to ESCRT assembly for lysosome repair. Nature 656, 216–226 (2026). 10.1038/s41586-026-10604-6

9 Tan, J. X. & Finkel, T. A phosphoinositide signalling pathway mediates rapid lysosomal repair. Nature 609, 815–821 (2022). 10.1038/s41586-022-05164-4

10 Wang, X. et al. The bridge-like lipid transport protein VPS13C/PARK23 mediates ER-lysosome contacts following lysosome damage. Nat Cell Biol 27, 776–789 (2025). 10.1038/s41556-025-01653-6

11 Lesage, S. et al. Loss of VPS13C Function in Autosomal-Recessive Parkinsonism Causes Mitochondrial Dysfunction and Increases PINK1/Parkin-Dependent Mitophagy. Am J Hum Genet 98, 500–513 (2016). 10.1016/j.ajhg.2016.01.014

12 Radulovic, M. et al. DFCP1-containing ER microdomains mediate lysosomal membrane repair. Nat Cell Biol (2026). 10.1038/s41556-026-02062-z

13 Axe, E. L. et al. Autophagosome formation from membrane compartments enriched in phosphatidylinositol 3-phosphate and dynamically connected to the endoplasmic reticulum. J Cell Biol 182, 685–701 (2008). 10.1083/jcb.200803137

14 Reinisch, K. M., De Camilli, P. & Melia, T. J. Lipid Dynamics at Membrane Contact Sites. Annu Rev Biochem 94, 479–502 (2025). 10.1146/annurev-biochem-083024-122821

15 Nahse, V. et al. ATPase activity of DFCP1 controls selective autophagy. Nat Commun 14, 4051 (2023). 10.1038/s41467-023-39641-9

16 Lukmantara, I. et al. PI(3)P and DFCP1 regulate the biogenesis of lipid droplets. Mol Biol Cell 33, ar131 (2022). 10.1091/mbc.E22-07-0279

17 Ismail, V. A., Naismith, T. & Kast, D. J. The NTPase activity of the double FYVE domain-containing protein 1 regulates lipid droplet metabolism. J Biol Chem 299, 102830 (2023). 10.1016/j.jbc.2022.102830

18 Ricciardi, S. et al. The role of NSP6 in the biogenesis of the SARS-CoV-2 replication organelle. Nature 606, 761–768 (2022). 10.1038/s41586-022-04835-6

19 Misra, M. et al. Structural basis for the ATP-dependence of omegasome biogenesis by DFCP1. bioRxiv, 10.64898/62026.64805.64827.728090 (2026).

20 Misra, S. & Hurley, J. H. Crystal structure of a phosphatidylinositol 3-phosphate-specific membrane-targeting motif, the FYVE domain of Vps27p. Cell 97, 657–666 (1999). 10.1016/s0092-8674(00)80776-x

21 Hayakawa, A. et al. Structural basis for endosomal targeting by FYVE domains. J Biol Chem 279, 5958–5966 (2004). 10.1074/jbc.M310503200

22 Chappie, J. S. et al. A pseudoatomic model of the dynamin polymer identifies a hydrolysis-dependent powerstroke. Cell 147, 209–222 (2011). 10.1016/j.cell.2011.09.003

23 Nixon-Abell, J. et al. Increased spatiotemporal resolution reveals highly dynamic dense tubular matrices in the peripheral ER. Science 354 (2016). 10.1126/science.aaf3928

24 Hu, J., Prinz, W. A. & Rapoport, T. A. Weaving the web of ER tubules. Cell 147, 1226–1231 (2011). 10.1016/j.cell.2011.11.022

25 Nishimura, T. et al. Autophagosome formation is initiated at phosphatidylinositol synthase-enriched ER subdomains. EMBO J 36, 1719–1735 (2017). 10.15252/embj.201695189

26 Mastronarde, D. N. SerialEM: A Program for Automated Tilt Series Acquisition on Tecnai Microscopes Using Prediction of Specimen Position. Microscopy and Microanalysis 9, 1182–1183 (2003).

27 Zheng, S. Q. et al. MotionCor2: anisotropic correction of beam-induced motion for improved cryo-electron microscopy. Nat Methods 14, 331–332 (2017). 10.1038/nmeth.4193

28 Elferich, J., Kong, L., Zottig, X. & Grigorieff, N. CTFFIND5 provides improved insight into quality, tilt, and thickness of TEM samples. Elife 13 (2024). 10.7554/eLife.97227

29 Kremer, J. R., Mastronarde, D. N. & McIntosh, J. R. Computer visualization of three-dimensional image data using IMOD. J Struct Biol 116, 71–76 (1996). 10.1006/jsbi.1996.0013

30 Zheng, S. et al. AreTomo: An integrated software package for automated marker-free, motion-corrected cryo-electron tomographic alignment and reconstruction. J Struct Biol X 6, 100068 (2022). 10.1016/j.yjsbx.2022.100068

31 Liu, Y. T. et al. IsoNet2 determines cellular structures at submolecular resolution without averaging. bioRxiv (2025). 10.64898/2025.12.09.693325

32 Agulleiro, J. I. & Fernandez, J. J. Tomo3D 2.0--exploitation of advanced vector extensions (AVX) for 3D reconstruction. J Struct Biol 189, 147–152 (2015). 10.1016/j.jsb.2014.11.009

33 Selzer, G. J. et al. napari-imagej: ImageJ ecosystem access from napari. Nat Methods 20, 1443–1444 (2023). 10.1038/s41592-023-01990-0

34 Burt, A. et al. An image processing pipeline for electron cryo-tomography in RELION-5. FEBS Open Bio 14, 1788–1804 (2024). 10.1002/2211-5463.13873

35 Cruz-Leon, S. et al. High-confidence 3D template matching for cryo-electron tomography. Nat Commun 15, 3992 (2024). 10.1038/s41467-024-47839-8

36 Mai, K. K. K. & Kang, B. H. Semiautomatic Segmentation of Plant Golgi Stacks in Electron Tomograms Using 3dmod. Methods Mol Biol 1662, 97–104 (2017). 10.1007/978-1-4939-7262-3_8

37 Pettersen, E. F. et al. UCSF ChimeraX: Structure visualization for researchers, educators, and developers. Protein Sci 30, 70–82 (2021). 10.1002/pro.3943

38 Ermel, U. H., Arghittu, S. M. & Frangakis, A. S. ArtiaX: An electron tomography toolbox for the interactive handling of sub-tomograms in UCSF ChimeraX. Protein Sci 31, e4472 (2022). 10.1002/pro.4472

39 Punjani, A., Rubinstein, J. L., Fleet, D. J. & Brubaker, M. A. cryoSPARC: algorithms for rapid unsupervised cryo-EM structure determination. Nat Methods 14, 290–296 (2017). 10.1038/nmeth.4169

40 Jumper, J. et al. Highly accurate protein structure prediction with AlphaFold. Nature 596, 583–589 (2021). 10.1038/s41586-021-03819-2

41 Casanal, A., Lohkamp, B. & Emsley, P. Current developments in Coot for macromolecular model building of Electron Cryo-microscopy and Crystallographic Data. Protein Sci 29, 1069–1078 (2020). 10.1002/pro.3791

42 Croll, T. I. ISOLDE: a physically realistic environment for model building into low-resolution electron-density maps. Acta Crystallogr D Struct Biol 74, 519–530 (2018). 10.1107/S2059798318002425

43 Afonine, P. V. et al. Real-space refinement in PHENIX for cryo-EM and crystallography. Acta Crystallogr D Struct Biol 74, 531–544 (2018). 10.1107/S2059798318006551

44 Prisant, M. G., Williams, C. J., Chen, V. B., Richardson, J. S. & Richardson, D. C. New tools in MolProbity validation: CaBLAM for CryoEM backbone, UnDowser to rethink “waters,” and NGL Viewer to recapture online 3D graphics. Protein Sci 29, 315–329 (2020). 10.1002/pro.3786

45 Barad, B. A. et al. EMRinger: side chain-directed model and map validation for 3D cryo-electron microscopy. Nat Methods 12, 943–946 (2015). 10.1038/nmeth.3541

46 Souza, P. C. T. et al. Martini 3: a general purpose force field for coarse-grained molecular dynamics. Nat Methods 18, 382–388 (2021). 10.1038/s41592-021-01098-3

47 Abraham, M. J. et al. GROMACS: High performance molecular simulations through multi-level parallelism from laptops to supercomputers. *SoftwareX* 1-2, 19–25 (2015). 10.1016/j.softx.2015.06.001

48 Orabi, E. A., Ozturk, T. N., Bernhardt, N. & Faraldo-Gomez, J. D. Corrections in the CHARMM36 Parametrization of Chloride Interactions with Proteins, Lipids, and Alkali Cations, and Extension to Other Halide Anions. J Chem Theory Comput 17, 6240–6261 (2021). 10.1021/acs.jctc.1c00550

49 Jorgensen, W. L., Chadrasekhar, J., Madura, J. D., Impey, R. W. & Klein, M. L. Comparison of simple potential functions for simulating liquid water. The Journal of Chemical Physics 79, 926–935 (1983).

50 Kroon, P. C. et al. Martinize2 and Vermouth provide a unified framework for molecular topology generation. Elife 12 (2025). 10.7554/eLife.90627

51 Kabsch, W. & Sander, C. Dictionary of protein secondary structure: pattern recognition of hydrogen-bonded and geometrical features. Biopolymers 22, 2577–2637 (1983). 10.1002/bip.360221211

52 Wassenaar, T. A., Ingolfsson, H. I., Bockmann, R. A., Tieleman, D. P. & Marrink, S. J. Computational Lipidomics with insane: A Versatile Tool for Generating Custom Membranes for Molecular Simulations. J Chem Theory Comput 11, 2144–2155 (2015). 10.1021/acs.jctc.5b00209

53 Boyd, K. J. & May, E. R. BUMPy: A Model-Independent Tool for Constructing Lipid Bilayers of Varying Curvature and Composition. J Chem Theory Comput 14, 6642–6652 (2018). 10.1021/acs.jctc.8b00765

54 Vogele, M., Kofinger, J. & Hummer, G. Molecular dynamics simulations of carbon nanotube porins in lipid bilayers. Faraday Discuss 209, 341–358 (2018). 10.1039/c8fd00011e

55 Shrestha, R. et al. Martini 3 Coarse-Grained Models for Carbon Nanomaterials. J Chem Theory Comput 21, 9035–9053 (2025). 10.1021/acs.jctc.5c00923

56 Tironi, I. G., Sperb, R., Smith, P. E. & van Gunsteren, W. F. A generalized reaction field method for molecular dynamics simulations. The Journal of Chemical Physics 102, 5451–5459 (1995).

57 Hess, B., Bekker, H., Berendsen, H. J. C. & Fraaije, J. G. E. M. LINCS: A linear constraint solver for molecular simulations. Journal of Computational Chemistry 18, 1463–1472 (1997). 10.1002/(SICI)1096-987X(199709)18:12<1463::AID-JCC4>3.0.CO;2-H

58 Bussi, G., Donadio, D. & Parrinello, M. Canonical sampling through velocity rescaling. J Chem Phys 126, 014101 (2007). 10.1063/1.2408420

59 Berendsen, H. J. C., Postma, J. P. M., van Gunsteren, W. F., DiNola, A. & Haak, J. R. Molecular dynamics with coupling to an external bath. The Journal of Chemical Physics 81, 3684–3690 (1984).

60 Bernetti, M. & Bussi, G. Pressure control using stochastic cell rescaling. J Chem Phys 153, 114107 (2020). 10.1063/5.0020514

61 Humphrey, W., Dalke, A. & Schulten, K. VMD: visual molecular dynamics. J Mol Graph 14, 33–38, 27–38 (1996). 10.1016/0263-7855(96)00018-5

62 Michaud-Agrawal, N., Denning, E. J., Woolf, T. B. & Beckstein, O. MDAnalysis: a toolkit for the analysis of molecular dynamics simulations. J Comput Chem 32, 2319–2327 (2011). 10.1002/jcc.21787

63 Buchete, N. V. & Hummer, G. Coarse master equations for peptide folding dynamics. J Phys Chem B 112, 6057–6069 (2008). 10.1021/jp0761665

64 Taubin, G. Estimation of planar curves, surfaces, and nonplanar space curves defined by implicit equations with applications to edge and range image segmentation. IEEE Transactions on Pattern Analysis and Machine Intelligence 13, 1115–1138 (1991). 10.1109/34.103273

65 Campeau, E. et al. A versatile viral system for expression and depletion of proteins in mammalian cells. PLoS One 4, e6529 (2009). 10.1371/journal.pone.0006529

